# p53 mutation in normal esophagus promotes multiple stages of carcinogenesis but is constrained by clonal competition

**DOI:** 10.1101/2022.02.03.478951

**Authors:** Kasumi Murai, Stefan Dentro, Swee Hoe Ong, Roshan Sood, David Fernandez-Antoran, Albert Herms, Vasiliki Kostiou, Benjamin A Hall, Moritz Gerstung, Philip H Jones

## Abstract

Aging normal human epithelia, such as the esophagus, accumulate a substantial burden of *TP53* mutant clones. These are the origin of most esophageal squamous carcinomas, in which biallelic *TP53* disruption is almost frequent. However, the cellular mechanisms by which *p53* mutants colonize the esophagus and participate in the subsequent stages of transformation are unclear. Here we show that inducing the *p53^R245W^* mutant in single esophageal progenitor cells in transgenic mice confers a proliferative advantage that drives clonal expansion but does not disrupt normal epithelial structure or function. Loss of the remaining *p53* allele in mutant cells does not increase their competitive fitness, creating a bottleneck to the development of chromosomally unstable *p53^R245W/null^* epithelium. In carcinogenesis, *p53* mutation does not initiate tumor formation, but tumors developing from areas with *p53* mutation and LOH are larger and show extensive chromosomal instability compared to lesions arising in wild type epithelium. We conclude that *p53* has distinct functions at different stages of carcinogenesis and that LOH within *p53* mutant clones in normal epithelium is a critical step in malignant transformation.

## Introduction

As humans age, our normal tissues become populated by mutant clones under strong positive selection, including genes recurrently mutated in cancer^1–9^. A typical example is the progressive increase in prevalence of clones carrying heterozygous *TP53* mutants in normal esophagus, which reaches 5-10% of the tissue by middle age rising to 15-30% for those in their 70s^2, 5^. This suggests that *TP53* mutants confer a competitive advantage on the cells that carry them^10^. A small subset of these clones in older people undergo loss of heterozygosity (LOH)^2^. Despite the accumulation of *TP53* mutants, human esophagus remains histologically normal and functionally intact, and genome stability is maintained other than frequent LOH at the *NOTCH1* locus^2^. Indeed, given the prevalence of *TP53* mutations the incidence of cancer seems surprisingly low.

In contrast to normal tissue, sequencing of squamous cell carcinoma (SCC) shows that the majority carry protein altering *TP53* mutations, most of which are associated with LOH^11, 12^(**Fig. 1a**). Multiregional sequencing indicates biallelic *TP53* disruption is a truncal event in SCC^11^. SCC are characterized by marked genomic instability, consistent with prior *TP53* loss being permissive of the survival of whole chromosomal aneuploid cells^12, 13^. Taken together, these results suggest *TP53* loss is required for SCC development and that tumors may originate via the rare clones with biallelic *TP53* disruption among the heterozygous *TP53* mutant population in normal tissue^2^.

**Fig. 1:**
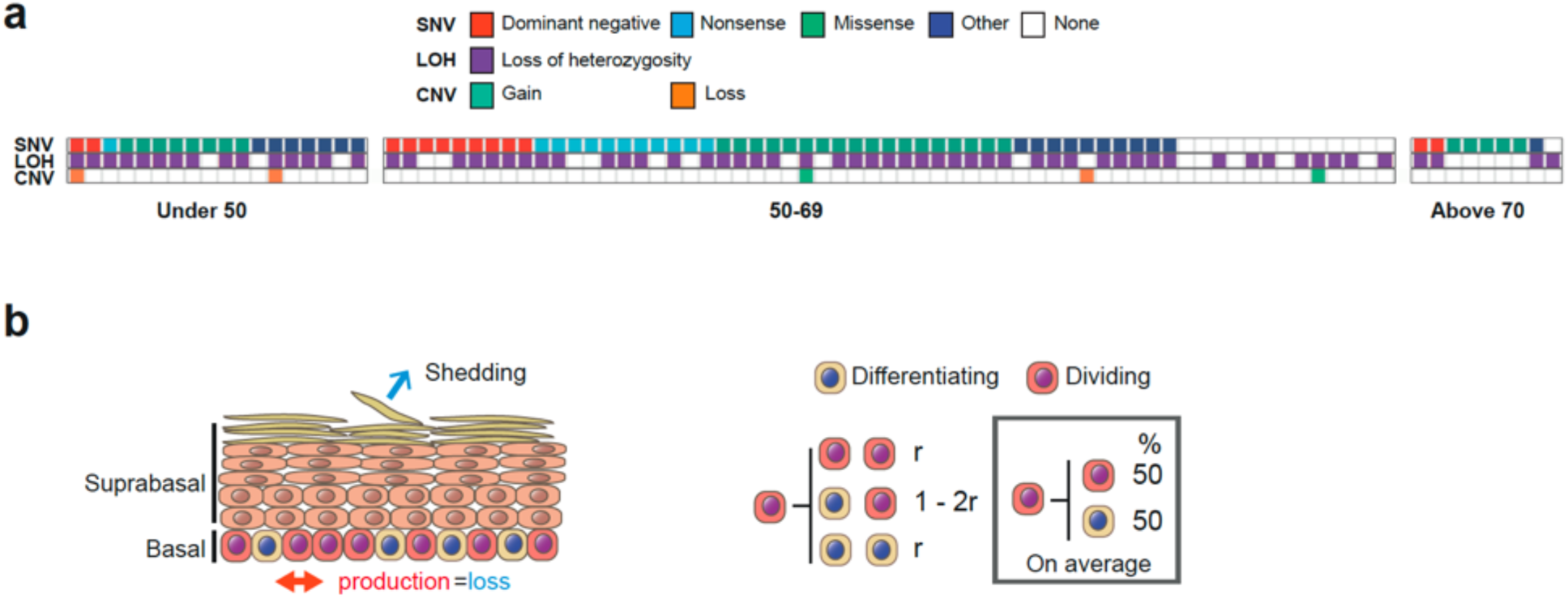
*TP53* mutation in Esophageal Squamous Cell Carcinoma (ESCC). **a**, 88 TCGA ESCC samples were analysed to identify the prevalence of genome alterations (single nucleotide variants, SNV, loss of heterozygosity, LOH & copy number variation, CNV) in the *TP53* gene. 74% of samples reported LOH over *TP53* and 84% reported SNVs. 64% of samples reported both SNVs and LOH over *TP53*. See **Supplementary Table 1** for source data. **b,** Cell behaviour in basal layer of normal homeostatic epithelium. On division, a progenitor may generate two dividing progenitors, two differentiating daughters, or one cell of each type. r is the probability of symmetric division outcome. On average, equal proportions of progenitor and differentiating cells are generated.

These observations motivated us to investigate the roles of *Trp53 (p53*) mutation on cell dynamics and clonal fitness in esophageal carcinogenesis using a transgenic mouse model.

Normal mouse esophagus has a simple, uniform structure that makes it ideal for studying the effects of mutations on cell behavior. The tissue consists of layers of keratinocytes. Dividing cells lie in the deepest, basal cell layer and are a single population of equipotential progenitor cells. Progenitor divisions generate either two progenitor daughters, two differentiating cells or one cell of each type (**Fig. 1b**). The likelihood of each division outcome is balanced so that one progenitor and one differentiating daughter are produced from an average division. It follows that equal numbers of progenitor and differentiating cells are generated across the progenitor population, thereby maintaining tissue homeostasis. Differentiating cells exit the cell cycle and leave the basal layer migrating to the tissue surface from which they are shed (**Fig. 1b**). Cells are continuously shed and replaced throughout life.

In transgenic mice it is possible to track the fate of cohorts of mutant cells in adult animals by lineage tracing. Scattered single progenitor cells are induced to express a mutation and a fluorescent reporter protein by genetic recombination. The daughter cells of the mutant progenitor may be visualized by 3D imaging and quantifying the proliferating and differentiating cells in clones provides insight into wild type and mutant cell dynamics^14–16^. Here we applied this approach to study how heterozygous *p53* mutant clones colonize the normal esophagus, the effects of loss of the second *p53* allele on clonal fitness and genome stability, and the role(s) of *p53* mutants in tumor development.

## Results

### Heterozygous *p53* mutation drives mutant clone expansion

We began by investigating whether the acquisition of a heterozygous *p53* mutation alters the behavior of esophageal progenitors using a transgenic mouse model. In *Ahcre^ERT^p53^flR245W-GFP/wt^* mice, hereafter termed *p53^*/^*^wt^, a cancer associated mutant allele of *p53*, *Trp53^R245W^*, corresponding to human *TP53^R248W^,* can be inducibly expressed following a genetic recombination event triggered by a drug regulated form of *cre* recombinase (**Fig. 2a**)^17, 18^. The *p53\** mutant is functionally different from a null allele and is frequently detected in squamous and other cancers ^19, 20^. Mutant cells can be tracked as expression of the mutant is linked to a green fluorescent protein (GFP) reporter. As a control, we tracked the behavior of *p53^wt/wt^* progenitors in the *Ahcre^ERT^Rosa26^flYFFP/wt^* (abbreviated to R^YFP^) mouse strain, where wild type cells are genetically labelled with a functionally neutral yellow fluorescent protein (YFP) reporter^15^.

**Fig. 2:**
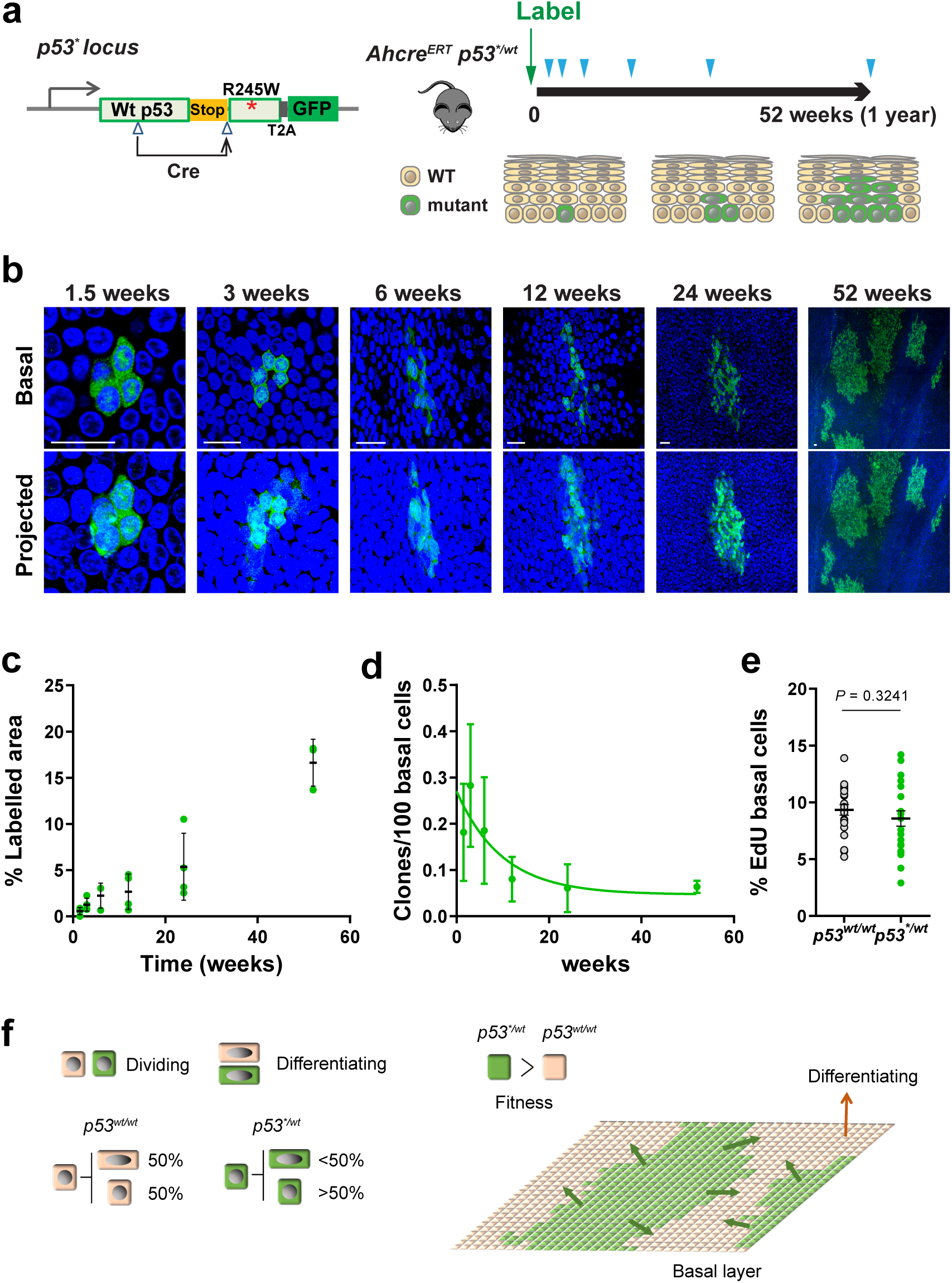
Heterozygous *p53^R245W^* (*p53^*/wt^*) mutant cell fate in mouse esophagus. **a,** Protocol and schematic of genetic lineage tracing in *Ahcre^ERT^ p53 ^flR245W-GFP/wt^* (*p53^*/wt^*) mice. Expression of the *p53* mutant allele and GFP reporter was induced in scattered single cells (Labelling) and esophagus samples were taken at indicated time points (triangles). The fate of mutant clones was examined by tracking the expression of GFP. n=4 mice per time point except n=3 at the 6-week and 52-week time points. **b,** Rendered confocal z stacks showing typical *p53^*/wt^* clones in esophageal epithelial wholemounts. Basal, top-down view of basal layer, Projected, top-down view through all nucleated cell layers. Green, GFP; blue, DAPI; Scale bars, 20µm **c,** Proportion of projected area labelled with GFP at indicated time points. Average value from 6 fields per animal. Error bars are mean ± s.e.m. **d,** Density of *p53^*/wt^* clones over the time. Error bars are mean ± s.e.m. **e**, Average percentage of EdU-positive basal cells in *p53^*/wt^* clones compared to non-GFP labelled (*p53^wt/wt^*) basal cells in the same mouse. EdU was administered an hour before sampling. n=22 mice across all time points. Error bars are mean ± s.e.m. *P* value, two-tailed paired Student’s t test. See **Supplementary Table 4-6**. **f,** Schematic illustration of *p53^*/wt^* cell behaviour. On average, wild type progenitors (*p53^wt/wt^*) produce equal proportions of dividing and differentiating cells across the population, whereas *p53^*/wt^* cells generate more dividing cells. This fate imbalance allows *p53^*/wt^* cells to outcompete wild type cells and expand in the tissue.

Heterozygous *p53\** or control YFP expression was induced in scattered single cells following the induction of *cre* recombinase in a cohort of adult mice. Expression of the GFP or YFP reporter is inherited by the progeny of the labelled cell, resulting in the formation of clones (**Fig. 2a and Extended Data Fig. 1a**). We prepared wholemounts of esophageal epithelium at different time points after induction, immunostained for the reporter protein and then performed 3D imaging of the tissue at single cell resolution using confocal microscopy. The number and location of cells in an unselected sample of clones was determined at each time point.

In *p53^wt/wt^* , R^YFP^ mice, the number of clones declined with time. This is due to differentiation, as if all the proliferating cells in a clone differentiate, the clone leaves the basal cell layer, migrates to the epithelial surface and is shed from the tissue^15^. However, the size of the remaining clones rose progressively, with the net result that the proportion of labelled cells in the epithelium remained constant over time. Such behavior is indicative of neutral competition between labelled and unlabeled wild type functionally equivalent progenitor cells (**Extended Data Fig. 1b,c**)^15^.

By contrast, in induced *p53^*/wt^* mice the area of labelled epithelium rose progressively indicating the mutant population had a competitive advantage over wild type neighbors (**Fig. 2b,c**). We found that GFP expression diminished in the upper suprabasal layers close to the epithelial surface, reflecting decreased transcription at the *p53* locus in differentiating cells^18^. To visualize mutant clones in all layers of the epithelium, we crossed *Ahcre^ERT^p53^*/wt^* mice with the *Rosa26^Confetti/wt^* reporter strain and induced recombination in single progenitors. This generated clones expressing red fluorescent protein from the constitutive *Rosa26* locus as well as GFP, and showed that differentiating *p53^*/wt^* cells extend through the upper differentiating layers of the tissue and reach the surface of epithelium, confirming mutant cells can be lost through shedding (**Extended Data Fig. 1d,e**)^18, 21^.

We next investigated the cellular mechanism(s) of *p53^*/wt^* clonal expansion. The mean number of basal layer cells/clone was consistently higher in *p53^*/wt^* clones than in YFP expressing clones in R^YFP^ mice at the corresponding time point (p<0.001 at 3, 6 and 12 weeks, two-tailed Mann-Whitney test, **Extended Data Fig. 1b**). The number of labelled mutant clones decreased over time consistent with mutant clone loss through differentiation (**Fig. 2d**). Further, the proportion of basal cells in S phase of the cell cycle labelled with a pulse of 5-ethynyl-2’-deoxyuridine (EdU) was similar in mutant and wild type cells, suggesting that the rate of cell division and proportion of proliferating cells were not substantially altered by the mutation (**Fig. 2e**).

Taken together, our observations are consistent with a model in which *p53^*/wt^* progenitors generate more progenitor than differentiating daughters per average cell division, so that the population grows despite no change in the rate of mutant cell division from that of wild type cells. To test this hypothesis we explored a simple quantitative simulation of clonal dynamics, a two-dimensional stochastic cellular automaton (CA) model (**Extended Data Fig. 2**). Mutant cells had a bias in cell fate, with an increased likelihood of dividing to generate two progenitor daughters compared with wild type cells. This proliferative advantage decreased when local cell crowding occurred, reflecting the spatial constraints of normal epithelium. This model gave a good quantitative fit to the observed clone sizes *in vivo*.

To gain further insight into the phenotype of *p53^*/wt^* epithelium, we induced *Ahcre^ERT^p53^*/wt^* mice at a high level and aged them for a year, by which time almost 70% of the epithelium was colonized by *p53^*/wt^* cells (**Extended Data Fig. 3**). The tissue remained macroscopically normal with no visible tumors. There was no apparent difference in histology or immunostaining for the marker of basal cells, KRT14 and the protein LOR, expressed in differentiating cells, comparing *p53^*/wt^* and wild type epithelium (**Extended Data Fig. 3**). Thus, as in humans, despite the presence of heterozygous *p53* mutants the esophagus remains functional and tissue integrity is maintained.

We conclude that cells carrying a heterozygous missense *p53* mutation colonize normal epithelium due to a subtle bias in progenitor fate towards proliferation (**Fig. 2f**). This results in a slowly expanding mutant population that will persist long term in the tissue, with the potential to undergo further genomic alterations.

### Loss of the second *p53* allele does not increase the competitive fitness of *p53\** mutant cells

The fitness of some heterozygous mutant genes that colonize the normal esophagus, such as *Notch1*, is increased following loss of the second allele, enhancing colonization and creating a selecting pressure towards LOH ^22^. To determine whether this was the case with *p53^*/wt^* cells we turned to a competition assay in organotypic culture (**Fig. 3a**) ^17^. Wild type cells were co-cultured for 4 weeks with either *p53^*/wt^* or *p53^*/-^* cells. Both mutant genotypes outcompeted WT cells to a similar degree. This indicates that *p53\** acts as a dominant negative for competitive fitness and hence that LOH confers no additional fitness advantage on *p53^*/wt^* cells (**Fig. 3b,c**). This observation has important implications, as if *p53^*/wt^* cells develop LOH they will compete neutrally with their *p53^*/wt^* neighbors, and the majority of the biallelic mutant clones may be lost through neutral drift rather than persisting to acquire further genomic alterations.

**Fig. 3:**
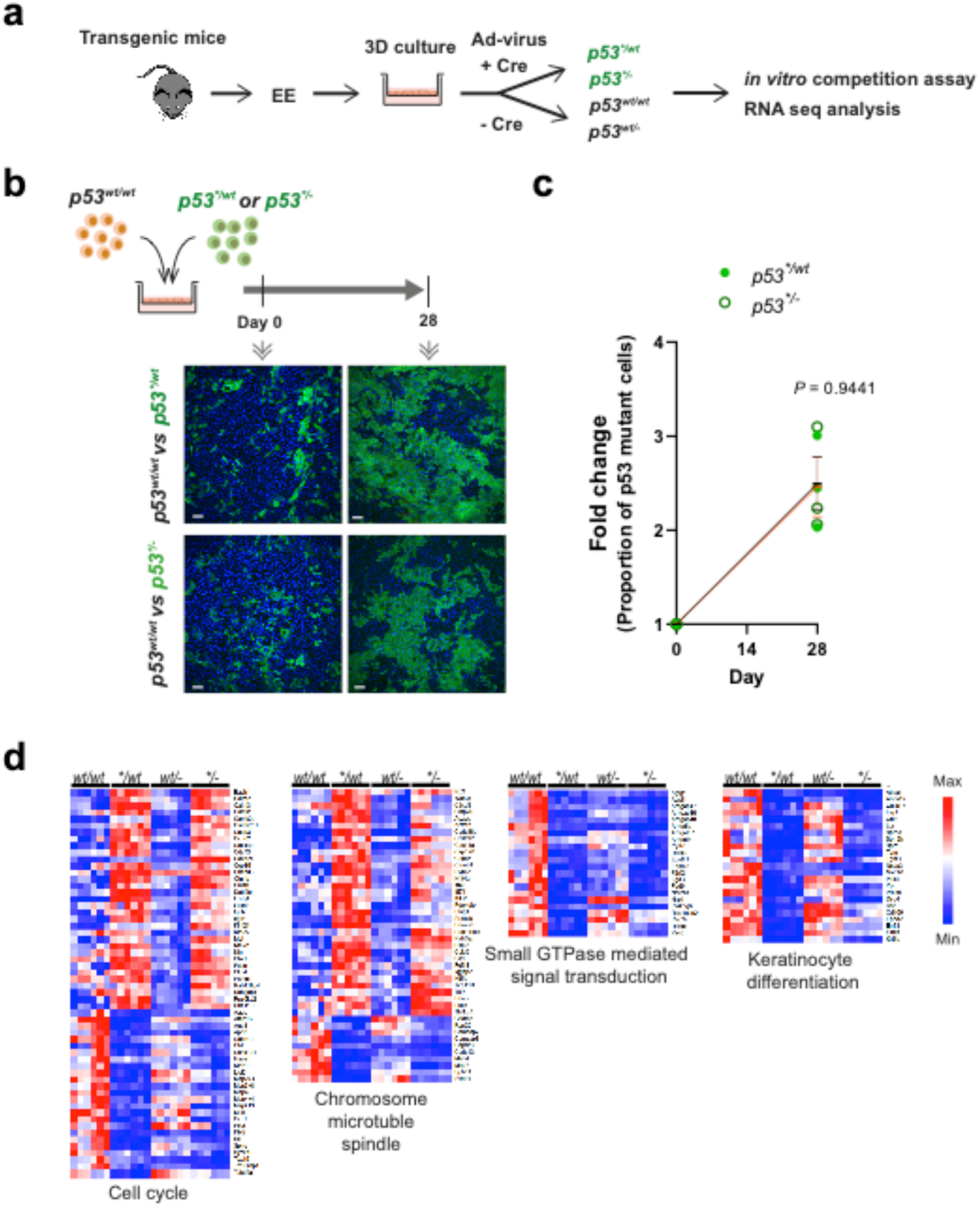
Effect of loss of heterozygosity (LOH) on mutant cell behaviour. **a,** A 3D primary culture system was used to characterise *p53* mutant cells *in vitro*. Esophageal keratinocytes were isolated from transgenic mice and *p53\** mutation was induced by adenovirus carrying cre recombinase. **b,** Cell competition assay. *p53^*/wt^* and *p53^*/-^* cells were co-cultured with *p53^wt/wt^* cells respectively and relative fitness was examined. Representative immunofluorescence images from three biological replicates at day 0 and 28 are shown. Green, GFP; blue, DAPI. Scale bars, 50 µm. **c,** Quantitation of cell competition assay by flow cytometry. Graph shows the fold change of proportion of GFP+ *p53^*/wt^* or *p53^*/-^* cell in the culture. Black (*p53^*/wt^*) and red (*p53^*/-^*) lines indicate mean and s.e.m. *P* value, two-tailed Student’s t-test, n= 3 replicate cultures. **d,** RNAseq analysis showing differentially expressed transcripts in *p53* mutant cells (*p53^*/wt^* and *p53^*/-^* compared to *p53^wt/wt^* or *p53^wt/-^*). Heatmaps were generated for genes associated with the cell cycle, cell division (chromosome, microtubule and spindle), small GTPase mediated signal transduction and keratinocyte differentiation. n=6 biological replicates per genotype. See **Supplementary Tables 7-9**.

To gain more insight into the basis of *p53^*/wt^* and *p53^*/-^* mutant fitness we analyzed the transcriptome in organotypic cultures of *p53^wt/wt^, p53^*/wt^*, *p53^wt/-^* and *p53^*/-^* esophageal epithelium. RNA sequencing identified transcripts differentially expressed in *p53^*/wt^* and *p53^*/-^* compared with *p53^wt/wt^* and *p53^wt/-^* cultures respectively (**Fig. 3a,d, Extended Data Fig. 4a,b and Supplementary Table 9**). The expression of multiple keratinocyte differentiation markers was down regulated and expression of cell cycle genes was altered in both *p53^*/wt^* and *p53^*/-^* compared to *p53^wt/wt^* and *p53^wt/-^* cells (**Fig. 3d and Supplementary Table 9**). Other mRNAs with altered expression included those encoding regulators of cytokinesis including microtubular, mitotic spindle and small GTPase mediated signal transduction genes. These results suggest that *p53\** mutant expression with or without a second wild type *p53* allele leads to mis-regulation of the cell cycle and decreases the differentiation of mutant cells, changes which parallel the tilt in cell fate from differentiation to proliferation observed *in vivo* in *p53^*/wt^* cells and explain the increased fitness of mutant cells.

### *p53^*/-^* epithelium

The genotype of SCC argues that they develop from the biallelic *p53* mutant clones that do succeed in persisting in the epithelium, so we set out to characterize the *p53^*/-^* phenotype. *Ahcre^ERT^p53^*/wt^* mice were crossed with a *p53^-/-^* strain. When the resulting *Ahcre^ERT^p53^*/-^* animals were induced, GFP expressing *p53^*/-^* clones competed in a background of unlabeled *p53^wt/-^* cells (**Fig. 4a,b**). The area occupied by *p53^*/-^* cells increased rapidly so that by a year almost the entire epithelium was replaced by *p53^*/-^* mutant cells, indicating they were substantially fitter than their *p53^wt/-^* neighbors. Consistent results were seen when *p53^*/-^* cells were co-cultured with *p53^wt/-^* cells (**Extended Data Fig.4c-f**).

**Fig. 4:**
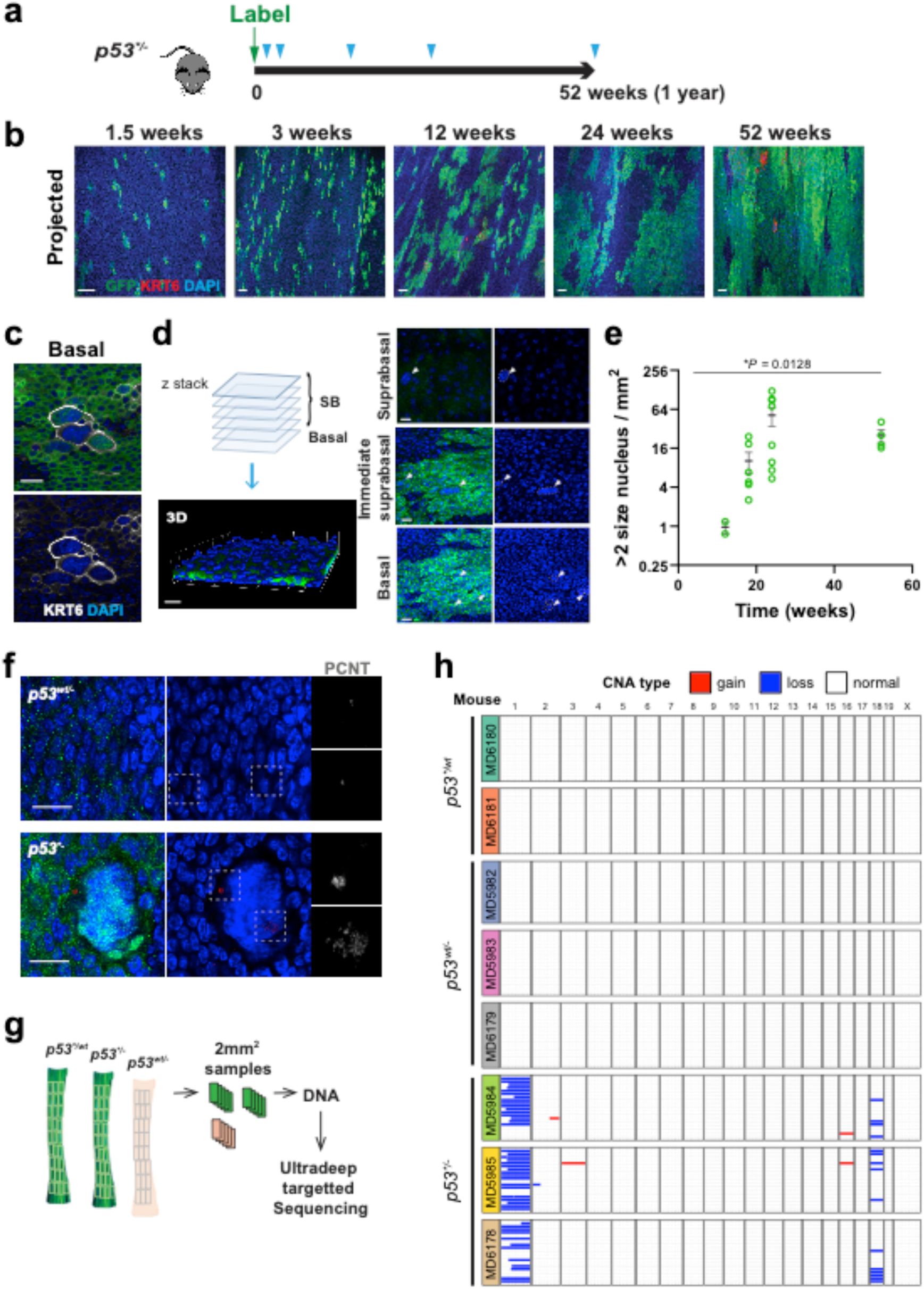
*p53^*/-^* esophageal epithelium. **a,** Protocol: *p53^*/-^* mice were generated with an inducible *p53\** allele and a *p53* null allele. Following induction, *p53^*/-^* clones competed in a *p53^wt/-^* background, and were sampled at indicated time points (triangles). **b,** Rendered confocal z stacks showing top down basal and projected views of *p53^*/-^* clones in wholemounts. Images are representative from n=2 mice for 1.5 and 3 week, n=4 for 12 week, n=7 for 24 week and n=4 for 52 week time points. Green, GFP; red, KRT6; blue, DAPI; Scale bars, 40µm. **c,** Confocal images showing cells with aneuploid appearance in *p53^*/-^*clone area. Green, GFP; grey, KRT6; blue, DAPI; Scale bars, 20µm. **d**, Confocal z stack images were used to reconstruct a 3D image of the wholemount sample and determine the z position of cells of interest. Aneuploid-like cells were found both in the basal layer and suprabasal layers. **e,** Number of cells with large (≥double size) nucleus in *p53^*/-^* clone area post induction. Error bars are mean ± s.e.m. *P* value determined by Welch’s ANOVA. n=2 mice for 12 week time point, n=6 for 18 week, n=8 for 24 week and n=4 for 52 week time point post induction. See Supplementary Table 11. **f,** Example images of above samples stained for PCNT. GFP-positive cells (green) are *p53^*/-^* and GFP negative cells are *p53^wt/-^* from the same EE. The yellow dashed boxes are shown at higher magnification in greyscale on the right. Aneuploid-appearing cells in the *p53^*/-^* clone area correlate with centrosome amplification. Red, PCNT; green, GFP; blue, DAPI. Scale bars, 20 µm. **g,** Wholemount epithelium was cut into a contiguous grid of 2mm^2^ pieces. DNA was extracted from each piece and subjected to ultradeep targeted sequencing. **h,** Summary of copy number analysis using targeted sequencing data, low coverage WGS of 2mm^2^ samples from same experiment is shown in **Extended Data Fig. 5f**.

In terms of phenotype, *p53^*/-^* epithelium looked macroscopically normal, tissue integrity was maintained and no tumors developed. Histology and the expression of KRT14 and LOR appeared unaltered compared with *p53^wt/wt^* and *p53^wt/-^* tissue (**Extended Data Fig.3**). However, confocal imaging of epithelial wholemounts revealed the progressive accumulation of scattered cells with enlarged nuclei in the GFP+ *p53^*/-^* areas but not the adjacent *p53^wt/-^* regions. These abnormal cells expressed the keratinocyte stress marker KRT6 and were detected both in the basal layer and the suprabasal layers (28.1% ± 4.2 in the suprabasal layer, n=6 from 24-52 week time point), arguing they can be lost by differentiation and shedding (**Fig. 4b-d**). The early accumulation of cells with large nuclei indicates these cells were initially generated at a faster rate than they were shed. At later time points, however, the number of these abnormal cells levelled off, indicating a balance between generation and shedding had been established (**Fig. 4e)**.

We speculated that the emergence of nuclear enlargement may indicate the development of genomic instability in the *p53^*/-^* epithelium. To investigate this, we first investigated the number of centrosomes in the cells with enlarged nuclei, as centrosomal amplification results in aneuploidy and tumorigenesis^23^. Staining for the centrosomal protein PCNT, pericentrin, revealed large foci consistent with centrosome amplification in the abnormal cells (**Fig. 4f)**. We concluded that in *p53^*/-^* epithelium, cells with centrosomal amplification develop and persist. Thus biallelic p53 disruption appears permissive of aneuploidy.

### Aging of *p53* mutant epithelium

In humans, *p53* mutant clones in normal epithelium may persist for many years. We therefore investigated the effects of aging on *p53\** mutant epithelium on the mutational landscape and copy number alterations in esophageal epithelium. Deep targeted sequencing for 142 genes associated with mouse and human epithelial cancers was performed on a gridded array of 2mm^2^ epithelial samples in esophagus 1 year after induction of *p53^*/wt^*, *p53^wt/-^* and *p53^*/-^* mutants (**Extended Data Fig. 5a, Methods and Supplementary tables 12, 13**)^24^. We identified 26 unique somatic coding mutations in a total area of 80mm^2^ *p53^*/wt^* epithelium (equivalent to 0.33 clones per mm^2^), and 50 and 46 mutations, each in total area 120mm^2^ epithelium (0.41 and 0.38 mutant clones per mm^2)^ in *p53^wt/-^* and *p53^*/-^* esophagus respectively (**Extended Data Fig. 5b**). The density of mutant clones in all three genotypes was similar to that in *p53^wt/wt^* esophagus (0.28 events per mm^2^)^24^. There was also no significant difference in the mutational burden between different genotypes (**Extended Data Fig. 5c**). The functional impact of the mutations was also similar in *p53\** mutant and wildtype esophagus (**Extended Data Fig. 5d**)^24^. Thus *p53\** mutation, with or without loss of the second *p53* allele, does not alter the incidence of age associated mutations in the normal mouse esophagus.

We then looked for evidence of copy number alterations (CNA) in the aged epithelium using off-target sequencing reads. In *p53^*/-^* tissue we observed recurrent loss of chromosome 1 and 18 in a small proportion of cells, consistent with clones with CNA within the samples (**Fig. 4g,h**). There was no evidence of such chromosomal instability in *p53^*/wt^* and *p53^wt/-^* epithelium. The calls were similar to those obtained from low coverage whole genome sequencing (**Extended Data Fig.5f**). These results argue that *p53* mutation with LOH leads chromosomal instability.

### Dynamics of *p53\** mutant clones in mutagenized epithelium

Aging human esophagus is a patchwork of mutant clones under strong competitive selection. To persist in such an environment *p53* mutant clones must outcompete neighboring clones, which expand until they encounter clones of similar fitness^24^. Mice treated with diethylnitrosamine (DEN), a well characterized mutagen found in tobacco smoke, develop a similar mutational landscape to older humans^24^. In addition, DEN treatment followed by aging leads to the development of esophageal tumors, both premalignant dysplasias and, more rarely, squamous cell carcinomas^24–26^.

We began by studying how *p53^*/wt^* mutant clones competed within a densely mutated normal epithelium. Mice were treated with DEN for 8 weeks, after which *p53\** expression was induced at a higher level than in a wild type background (**Fig. 2**) to allow for the elimination of *p53^*/wt^* clones by DEN induced mutants. The area occupied by *p53\** cells was monitored for up to a year post-induction (**Fig. 5a**). As expected, in controls which were not treated with DEN *p53*^*/wt^ mutant clones expanded, fused and occupied the majority of the epithelium over the year after induction (**Fig. 5b,c**). In contrast, the expansion of *p53^*/wt^* mutant clones was severely reduced in the DEN-treated esophageal epithelium, reaching only 5.5% of the area of the esophagus by at the 52 week time point.

**Fig. 5:**
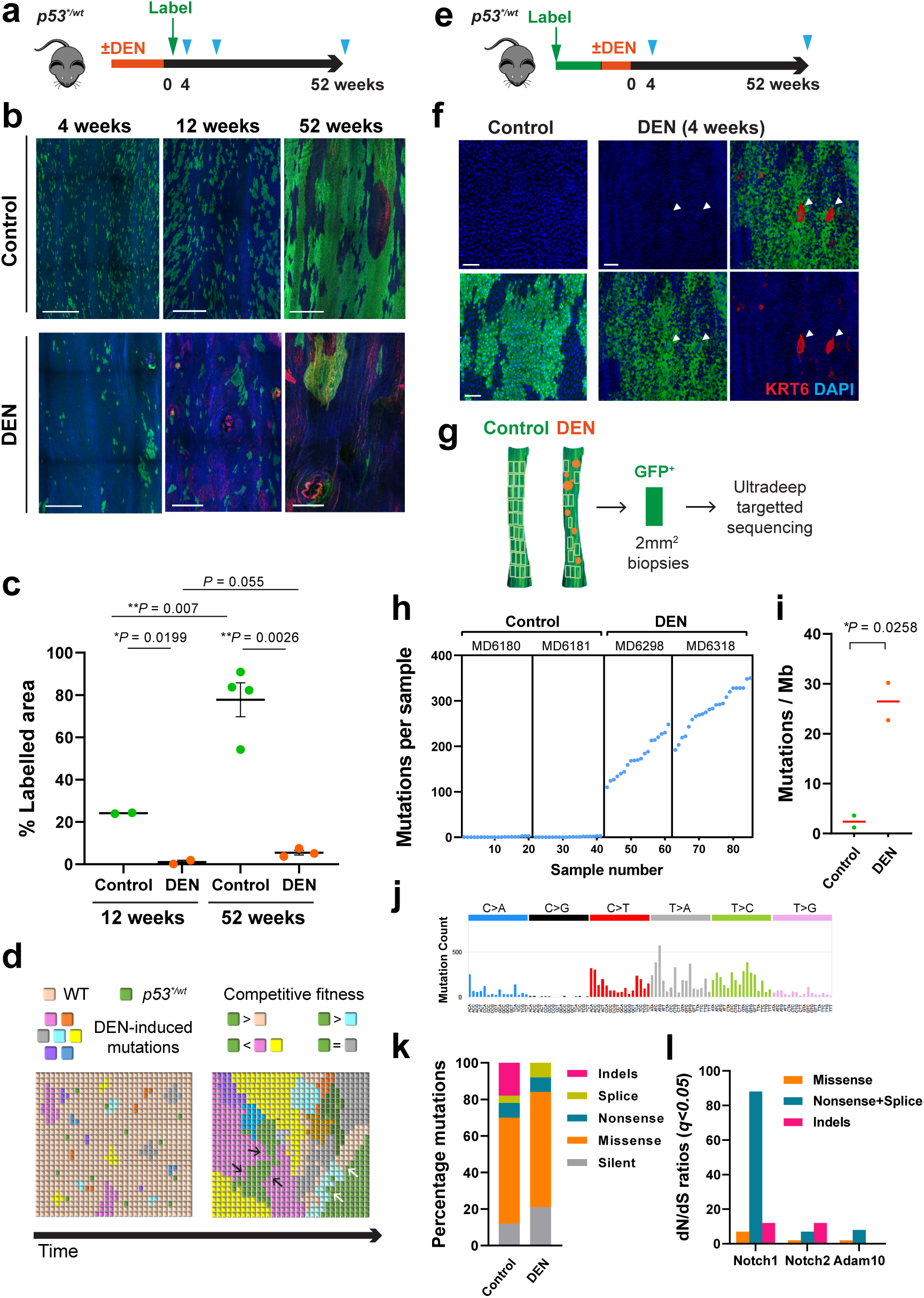
*p53^*/wt^* mutant clones in mutagenized epithelium. **a,** Protocol: *p53^*/wt^* mice were treated with for 8 weeks with DEN or vehicle followed by clonal induction, samples were collected at indicated time points (triangles). **b,** Confocal z stacks showing projected top down views of *p53^*/wt^* clones in epithelial wholemounts from control and DEN-treated mice. Representative images from two independent experiments. n=4 control and n=3 DEN-treated mice at 4-week time point, n=2 per group at 12-week time point, n=6 control and n=4 DEN-treated mice at 52-week time point. Green, GFP; Red, KRT6; blue, DAPI. Scale bars 500 µm. **c,** Average proportion of projected labelled area at indicated time points. Number of mice as in **b**. Error bars indicate mean ± s.e.m., p values were determined by Welch’s t test. See Supplemental Table 14. **d,** Schematics of *p53^*/wt^* clone behaviour in DEN-treated tissue. *p53^*/wt^* outcompetes wild type cells, but not all DEN-induced mutant cells. Many *p53^*/wt^* clones are lost by the 4-week time point. Surviving *p53^*/wt^* clone 1-year time vary widely in size and are found in both epithelium and tumors. *p53^*/wt^* clone can be outcompeted by surrounding fitter clones (black arrows) or itself be fitter than adjacent mutant clones (white arrows). **e,** Protocol: *p53^*/wt^* mice were treated with DEN or vehicle for 8 weeks or after colonisation by *p53^*/wt^* mutant cells. Samples were collected at indicated time points (triangles). **f,** Confocal images showing top down views of basal layer of wholemounts from *p53^*/wt^* induced mice 4 weeks after DEN treatment (n=9 mice) compared to control (vehicle, n=4 mice). Green, GFP; grey, CDH1; red, KRT6; blue, DAPI. White arrowheads indicate large nuclei observed in *p53^*/wt^* clone area following DEN treatment. Scale bar, 50µm. **g,** Sequencing analysis at 1 year time point (Fig. 5e) 2mm^2^ grid biopsies of epithelium and DNA extracted from each sample were subjected to ultradeep targeted sequencing. **h,** Number of mutations per sample. Every dot corresponds to a sample. n=2 mice per condition. **i,** Estimated mutation burden in untreated control and DEN-treated samples. p value by unpaired Student’s t-test. Red lines indicate mean value. **j,** Mutational spectrum of DEN-treated *p53^*/wt^* epithelium. **k,** Percentage of mutation types identified in each condition. **l,** Positively selected somatic mutations in DEN-treated samples. dN/dS ratios for missense, truncating (nonsense + splice) and indels. See **Supplementary Table 12-13**.

Both large and small or fragmented *p53^*/wt^* clones were seen, consistent with clonal competition^18, 24^ (**Fig. 5b**). Previous deep sequencing study identified numerous mutant clones in mouse esophagus treated with same dose of DEN^24^. Thus *p53^*/wt^* clonal expansion is restricted by competing DEN induced mutant clones in the epithelium (**Fig. 5d**) ^24^. However, whilst their expansion is constrained, *p53^*/wt^* mutant clones are sufficiently fit to persist in this highly competitive clonal landscape (**Fig. 5b**). These findings in mice recapitulate the restricted expansion and long-term persistence of heterozygous oncogenic *TP53* mutants in the highly competitive densely mutated normal epithelium of aging human esophagus ^2^.

Next, we performed an alternative protocol to model the effect of additional mutagenesis on *p53^*/wt^* cells, as occurs within persisting *p53* mutant clones in humans. *p53^*/wt^* mice were highly induced, the resulting clones allowed to expand, and the animals then treated DEN (**Fig. 5e**). Within 4 weeks after DEN treatment, cells expressing KRT6 with abnormally large nuclei were detected in GFP+ expressing areas, resembling those observed in *p53^*/-^* epithelium, changes consistent with the emergence of aneuploid cells (**Fig. 4 and 5f**).

To further examine the effect on mutational landscape, we performed deep-targeted sequencing of the epithelium. This revealed 102 mutant clones/mm^2^ compared to ∼2.4/mm^2^ in untreated *p53^*/wt^* mice **(Fig. 5g,h, Methods**). The mutational burden was ∼26 mutations per megabase in the DEN treated animals compared to ∼2.4 in untreated *p53^*/wt^* mice (**Fig. 5i**). Most mutations were protein altering and the mutation spectrum was dominated by T>A/A>T, T>C/A>G and C>T/G>A alterations (**Fig. 5j,k**), typical of DEN exposure^24^. To identify positively selected mutant genes in DEN- treated *p53^*/wt^* epithelium, we calculated the dN/dS (non-synonymous-to synonymous mutation) ratios across all sequenced genes. Mutant *Notch1*, *Notch2* and *Adam10* were positively selected in *p53^*/wt^* epithelium (**Fig. 5l**). These results are all very similar to those in DEN exposed wild type epithelium indicating that *p53\** mutation does not alter the mutational landscape.

Next, we explored the impact of the heterozygous *p53^*/wt^* mutation on tumor formation, in two different protocols (**Extended Data Fig. 6a**). In protocol 1, mice were treated with DEN *p53\** mutation induced and animals aged, modelling the emergence of *p53* mutant clones within the already mutated landscape (**Extended Data Fig. 6a, protocol 1**). Persistent GFP+ *p53^*/wt^* clones were detected both within tumors but also in normal areas of EE at a year post induction (**Fig. 5b**). The 20% of lesions that contained GFP+ *p53\** mutant cells were larger on average than the GFP negative tumors (**Extended Data Fig. 6b**). These observations argue the heterozygous *p53\** mutation does not initiate tumor formation but does promote tumor growth. In protocol 2, mice were induced and *p53\**clones allowed to expand, mice were then treated with DEN and aged, modelling the effect of additional mutations within *p53* mutant epithelium (**Extended Data Fig. 6a**). After 24 weeks, the number of tumors did not differ significantly between *p53^*/wt^* induced and uninduced control (*p53^wt/wt^*) animals in either protocol (**Extended Data Fig. 6c**). The timing of *p53^*/wt^* induction relative to DEN mutagenesis thus had no significant impact on the incidence of tumors. However, tumors originating from *p53^*/wt^* epithelium were significantly larger than those in the uninduced controls (**Extended Data Fig. 6a,d, protocol 2**).

We then went on to characterize the *p53^*/wt^* expressing lesions. Histology of macroscopic tumors showed the features of high-grade dysplasia, with loss of normal differentiation, architectural abnormalities, cell crowding and loss of polarity (14 out of 16 tumors showed dysplasia or SCC) (**Supplementary table 19**). In some cases, lesions extended deep into the submucosa (**Extended Data Fig. 6e**). Two out of eleven *p53^*/wt^* lesions developed into SCC by the 1 year time point (**Extended Data Fig. 6f-i**). Immunostaining of the SCC demonstrated a loss of expression of the terminal differentiation marker and cornified envelope precursor protein LOR. We also observed widespread expression of KRT14 which is normally confined to the basal layer and KRT6 which is a marker for murine premalignant and malignant esophageal tumors^25^. In addition, immune cell infiltrates were detected by staining for the pan leukocyte marker, CD45. Staining for PCNT revealed multiple or clustered foci indicative of centrosome amplification (average 42% cells with >2 centrosomes (**Extended Data Fig. 6j,k**). This is significantly more frequent than in tumors arising from *p53^wt/wt^* esophagus where such PCNT foci occur in an average 15% of cells.

In parallel, we also examined the mutational landscapes of 11 of the larger tumors. For 5 lesions the adjacent epithelium was also sequenced. Deep targeted sequencing of 142 genes, with a mean on-target depth of coverage 183x, identified a total of 1373 mutations from 11 tumors and 722 from 5 adjacent epithelium (**Extended Data Fig. 7a,b and Methods**). The mutation burden and functional impact of mutations was similar between tumors and epithelium and the mutational spectrum was typical of DEN exposure (**Extended Data Fig. 7c-e**). Analysis of the variant allele frequency (VAF) indicated the epithelium was polyclonal whereas tumors other than SCCs were monoclonal (**Extended Data Fig. 7f**). In addition, within tumors, mutant *Notch1* and *Atp2a2* were positively selected (**Extended Data Fig. 7g**)^27^. We also looked for evidence of CNA in *p53^*/wt^* tumors using off target reads of the above sequencing. 3 out of 12 macroscopic tumors and 2 out of 18 microscopic lesions had detectable CNA (**Extended Data Fig. 7h,i**), consistent with observation of a higher proportion of centrosome amplification in these tumors. This contrasts with DEN induced tumors from *p53^wt/wt^* mice, only 3% (2 of 64 tumors) exhibit CNA^27^.

### Effect of mutagenesis on *p53^*/-^* epithelium

Since the genotype of human esophageal SCC suggests they arise from *TP53* mutant cells with LOH, we evaluated the effect of mutagenesis and aging of *p53^*/-^*epithelium. *Ahcre^ERT^p53^*/-^* mice were induced to generate *p53^*/-^* clones. Animals were exposed to a lower dose of DEN than used above, to reduce the risk of malignancy in *p53^*/-^* cells recombined outside the esophagus, and then aged (**Fig. 6a**). At 1 year, the esophagus was mosaic, with unrecombined *p53^wt/-^* areas and *p53^*/-^* epithelium allowing a direct comparison of tumor burden in the same animal. This showed the number of tumors was significantly higher in GFP+ *p53^*/-^* area than *p53^wt/-^* area (**Fig. 6b,c**). Furthermore, when tumors and adjacent epithelium were sequenced to detect CNA, there was minimal CNA in *p53^wt/-^* epithelium, but extensive CNA in both *p53^*/-^* epithelium and the tumors within it (**Fig. 6d,e)**.

**Fig. 6.**
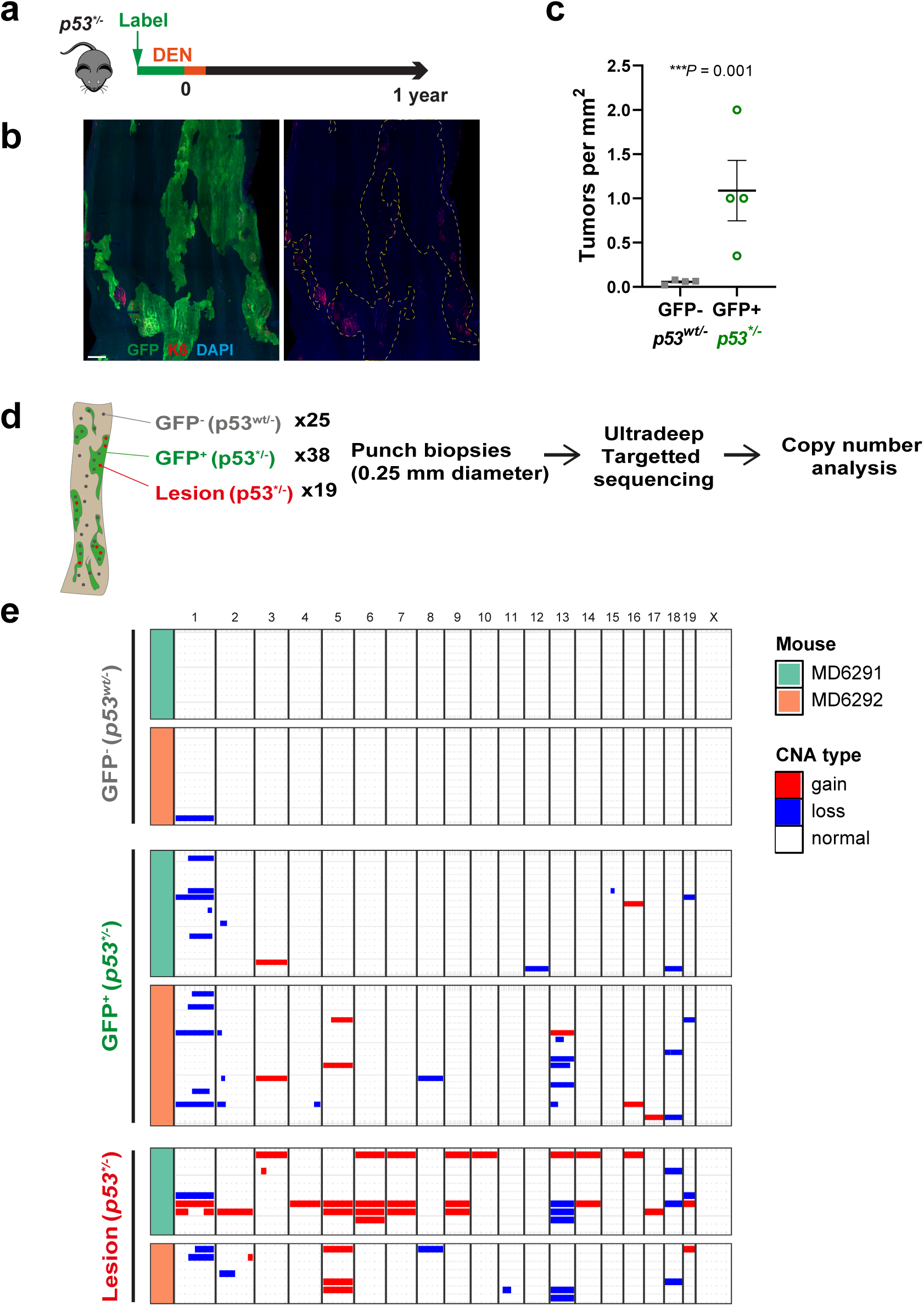
Effect of mutagenesis and aging *p53^*/-^* epithelium. **a,** Protocol: Induced *p53^*/-^* mice were treated with DEN for 2 weeks and aged**. b,** Confocal image of *p53^*/-^* induced epithelium at 1 year post DEN treatment. typical example from n=4 mice. Green, GFP; red, KRT6; blue, DAPI. Scale bar, 400 µm. **c,** Number of *p53^*/-^* (GFP+) and *p53^wt/-^* (GFP-) tumors. n=4 mice, p value was determined by two-tailed ratio paired t-test. **d,** Sequencing analysis of DEN treated *p53^*/-^* epithelium at 1 year time point. Micro- punch biopsies were taken from physiologically normal epithelium, GFP-negative *p53^wt/-^* or GFP- positive *p53^*/-^* clone area and from lesions (*p53^*/-^*). Chromosomal alterations were analysed using targeted sequencing data. **e,** Summary of copy number alterations. Gain and loss of chromosomes was predominantly detected in *p53^*/-^* clone area and lesions which arose from *p53^*/-^* cells. n=2 mice; 25 biopsies for *p53^wt/-^*, 38 biopsies for *p53^*/-^* clone areas, 19 biopsies for lesions. See Supplementary Table 23 and 24.

Taken together, above results show that the heterozygous *p53\** mutant promotes tumor growth, with a modest effect on CNA, whereas *p53\** combined with loss of the second p53 allele in the setting of low dose mutagen exposure increases the number of tumors and results in marked chromosomal instability both in normal epithelia and in tumors.

## Discussion

Our results establish that the *p53\** mutation has discrete roles at distinct stages of carcinogenesis. Heterozygous *p53\** progenitors acquire a proliferative advantage through a bias in cell fate, so that on average mutant cell divisions produce an excess of progenitors over differentiated cells. This fate imbalance is sufficient to drive clonal expansion in both wild type and mutagenized epithelium which contains competing clones of varying fitness^24^. A similar cellular mechanism is shared by diverse mutations that cause esophageal clonal expansion, including activating *Pik3ca^H1047R^*, *Notch1^wt/-^* and dominant negative mutant *Maml1*^14, 28, 29^. In common with these other mutants, after an initial phase of expansion *p53^*/wt^* has a minimal epithelial phenotype. Once mutant cells are surrounded by cells which are themselves mutant, cell fate bias decreases and reverts towards balance allowing the epithelium to retain its normal appearance and functional integrity^24^. Furthermore, we observed no increase in age associated mutation rate above that of wild type esophagus or any detectable CNA in *p53^*/wt^* epithelium. Heterozygous *p53\** induced at clonal density expands to establish a mutant population that persists long term within the esophagus. These findings are consistent with the slow accumulation of heterozygous *TP53* mutants over decades in normal human esophageal epithelium.

Despite the burden of *TP53* mutants reaching up 20% of cells in older humans, the life time risk of esophageal cancer is only about 1%^2, 5, 30^. This argues there may be bottlenecks that restrict transformation of mutant epithelium. 90% of esophageal dysplastic lesions as well as cancers exhibit disruption of both *TP53* alleles arguing there is strong selection of clones with inactivation of wild type allele in *TP53* heterozygous mutants during malignant transformation^11, 12, 31^. In normal epithelium, the cell competition experiments reported here indicate no change in fitness comparing *p53*^/-^* and *p53^*/wt^* cells. Similar observations have been made in the hematopoietic system^32^. It follows that when a cell within a heterozygous mutant clone loses the second allele it will compete neutrally with its neighbors and generate a clone that is likely to be lost from the tissue within a few rounds of cell division^15^. If replicated in human epithelium, this ‘neutral bottleneck’ would explain the scarcity of *TP53* mutant clones with LOH observed in normal epithelia, and would represent an important barrier against transformation^2^.

The impact of biallelic *TP53* loss not increasing fitness is well illustrated by comparing with mutant *NOTCH1*. Loss of the second allele of *Notch1* in mouse esophagus increases the fitness of heterozygous mutant clones substantially^29^. In humans most *NOTCH1* clones in normal epithelium have LOH of *NOTCH1* and spread to occupy the majority of the normal epithelium by middle age^2^. If *TP53* behaved in a similar manner, the proportion of epithelium qualified for tumor progression would be very much higher than is observed.

Once the ‘neutral bottle neck’ is overcome, the *p53^*/-^* epithelium exhibits evidence of genome instability in the form of giant multinucleated cells and clonal expansions carrying CNA not seen in esophagus that retains a wild type copy of *p53*. The scarcity of *TP53* mutant clones with LOH in humans makes it difficult to know if such a phenotype occurs prior to transformation in human esophagus, but marked CNA is an early feature of both premalignant lesions and SCC with biallelic *TP53* loss^11^. The suggestion that *p53^*/-^* cells are better qualified for transformation than *p53^*/wt^* seems borne out in mutagenesis studies. Tumor size was increased in p53^*/wt^ mutant epithelium, arguing the mutation promotes tumor growth, but there was no increase in the number of tumors and minimal CNA. By contrast, after mutagen treatment there was extensive CNA in both *p53^*/-^* epithelium and the tumors derived from it compared with *p53^*/wt^* tissue, and the *p53^*/-^* tumors were more numerous.

In conclusion these findings highlight how oncogenic *p53* mutations have distinct effects at different stages of transformation (**Fig. 7**). The promotion of clonal expansion results in minimal disruption to normal epithelium, indeed is tolerated in humans over decades. Heterozygous mutants have a low risk of transformation and do not contribute to genome instability. Once the second allele is lost, a clone that persists and escapes the ‘neutral bottleneck’ may develop additional mutations or CNA that promotes clonal expansion and/or tumor formation^33–35^. Within established tumors, *p53* mutation is advantageous, favoring tumor growth. The multiple facets of *p53* mutant biology must be placed in the context of clonal competition to understand epithelial carcinogenesis.

**Fig. 7.**
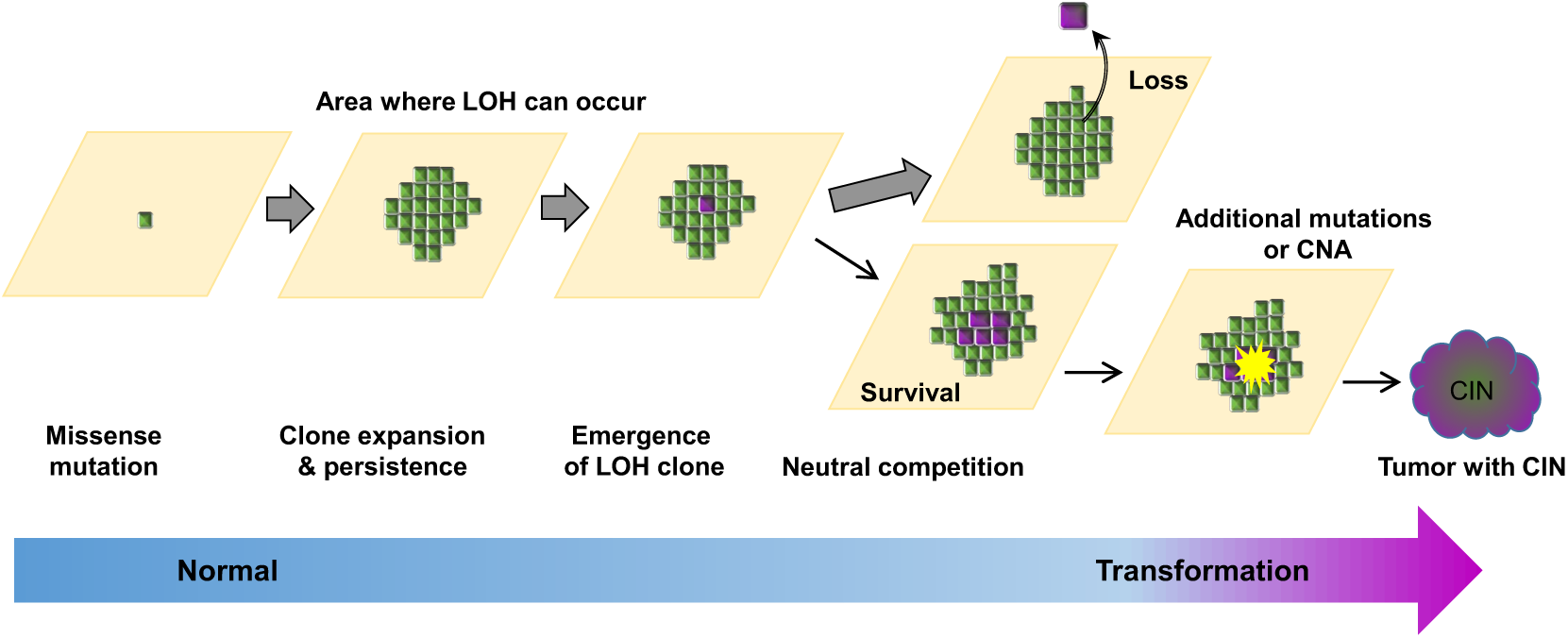
Mutant *p53* in normal epithelia in carcinogenesis. Heterozygous *p53* mutation in single cells in normal epithelia confers a proliferative advantage, clonal expansion and a population of mutant cells that persists to undergo LOH. Most cells with LOH are lost by neutral competition, but the few clones that expand and persist acquire CNA and may progress to form tumors. Once a tumor has formed, *p53* mutation enhances tumor growth.

## Acknowledgements

This work was supported by grants from the Wellcome Trust to the Wellcome Sanger Institute (098051 and 296194) and Cancer Research UK Programme Grants to P.H.J. (C609/A17257 and C609/A27326). B.A.H. was supported by the Medical Research Council (Grant-in-Aid to the MRC Cancer unit grant number MC_UU_12022/9 and NIRG to B.A.H. grant number MR/S000216/1). B.A.H. acknowledges support from the Royal Society (grant no. UF130039). S.D. benefited from the award of an ESPOD fellowship, 2018-21, from the Wellcome Sanger Institute and the European Bioinformatics Institute EMBL-EBI. We thank Esther Choolun, Tom Metcalf and staff at the MRC ARES and Sanger RSF facilities for excellent technical support and Michael Hall for statistical advice.

## Author Contributions

K.M., designed and performed experiments, assisted by D.F.A. and A.H. D.F.A performed the RNAseq experiments. R.S., S.H.O., S.D., analyzed sequence data. V.K. and B.A.H. performed clone simulations. K.M., S.D. B.A.H. and P.H.J. wrote the paper. B.A.H., M.G. and P.H.J. supervised the research.

## Materials and Methods

### Animals

Experiments were conducted according to UK government Home Office project licenses PPL22/2282, PPL70/7543 and PPL4639B40. Animals were housed in individually ventilated cages and fed on standard chow and maintained on a *C57/Bl6* genetic background, at specific and opportunistic pathogen free health status and were immune competent. Adult mice were used for in vivo experiments and no animals were involved in previous experiments and were drug naive prior to the start of experiments. Both male and female animals were used for experiments.

Ahcre^ERT^Rosa^flEYFP/wt^, Ahcre^ERT^Rosa26^flconfetti/wt^, and Ahcre^ERT^Trp53^flR245W-GFP/wt^ knock-in mice were generated as previously described. Ahcre^ERT^Trp53^flR245W-GFP/-^ mice were generated by crossing Ahcre^ERT^Trp53^flR245W-GFP/flR245W-GFP^ with heterozygous p53 knockout mice, p53^+/-^.

In the *Ah^creERT^* line, transcription from a transgenic *CYP1A1* (arylhydrocarbon receptor, *Ah*) promoter is normally tightly repressed (Kemp et al., 2004). Following treatment with the non-genotoxic xenobiotic β-napthoflavone the *Ah* promoter is induced and a *cre* recombinase-mutant oestrogen receptor fusion protein (creERT) is expressed. In the presence of tamoxifen, the creERT protein enters the nucleus to mediate recombination.

For lineage tracing experiments, the relevant floxed reporter line which was crossed onto the *Ah^creERT^* strain were induced by a single interaperitoneal (i.p) injection of 80 mg/kg β-napthoflavone (βNF)and 1 mg tamoxifen (Tam) at 11-16 weeks of age. For some experiments, in order to achieve optimal clone density, xx mg/kg βNF and xx mg Tam was used for for *Ahcre^ERT^Trp53^flR245W-GFP/wt^* and *Ahcre^ERT^Trp53^flR245W-GFP/-^* GFP and YFP expressing clones were visualised by immunostaining with an anti-GFP antibody.

### Mutagen treatment

To induce mutations in the oesophageal epithelium, mice were administered with DEN in sweetened drinking water (40 mg/L) for 3 days a week (Monday, Wednesday and Friday) for 8 weeks. Control mice received sweetened water as vehicle for the length of the treatment.

### Whole mount sample preparation

Mouse oesophagus was dissected, cut longitudinally and the muscle layer removed by pulling with forceps. Entire tissue was incubated for 2.5 hours in 5mM EDTA at 37°C before and then the epithelium was carefully separated from underlying submucosa using fine forceps followed by fixation in 4% paraformaldehyde for 30 min at room temperature. The tissues were then washed in PBS and stored at 4°C.

### Immunofluorescence

For staining, wholemounts were blocked in staining buffer (0.5% Bovine Serum Albumin, 0.25% Fish Skin Gelatin, and 0.5% Triton X-100 in PBS with 10% goat or donkey serum according to the secondary antibody used) for 1 hour at room temperature. Samples were incubated with primary antibody in staining buffer overnight, washed in PBS containing 0.2% Tween-20 four times, incubated with fluorchrome-conjugated secondary antibody for 2 hours at room temperature and washed as before After the final wash, samples were incubated with 40,6-diamidino-2-phenylindole (DAPI, 1 µg ml^-1^) or SiR-Hoechst (SiR-DNA) (Spirochrome, SC007) to stain cell nuclei and mounted on slides using Vectashield Mounting Medium (Vector Labs).

EdU incorporation was detected with a Click-iT imaging kit (Life Technologies) according to manufacturer’s instructions.

### Antibodies

Primary antibodies used in this study are as follows: anti-GFP/YFP (ThermoFisher Scientific, A10262), anti-Cytokeratin 14 (Covance, PRB-155P), anti-Cytokeratin 6 (Biolegend, 905701), anti-pericentrin (abcam, ab4448), anti-Aurora Kinase B (abcam, ab2254), anti-p53 (CM5)(Vector Laboratories, VP-P956), anti-αTubulin (Cell Signaling Technology, #3873), anti-CD45 (Biolegend, 103102), anti-CD31 (abcam, ab7388), anti-Loricrin (Covance PRB-145P), Alexa Fluor 647 anti-CD49f (Biolegend, 313610).

All secondary antibodies are Alexa Fluor conjugated: 488 anti-chicken (Jackson ImmunoResearch, 703-545-155), 555 anti-rabbit (ThermoFisher Scientific, A31572), 647 anti-mouse (ThermoFisher Scientific, A32787), 488 and 647 anti-rat (ThermoFisher Scientific, A21208 and A48272).

### Frozen sections and histology

Tissues were frozen in O.C.T and cut at 14 µm thickness in the cryostat. Sections were then fixed in 4% paraformaldehyde in PBS for 10 minutes and washed in PBS before staining with hematoxylin and eosin or immunostaining as described above.

### Pathology of macroscopic tumors

Pathological assessment of tumors was carried out by MRC Metabolic Diseases Unit [MC_UU_00014/5].

### Imaging

Confocal images were acquired on Leica TCS SP5 II or SP8 microscopes using 10x, 20x, 40x or 64x objectives. Typical settings for acquisition of z stacks were optimal pinhole, line average 4 scan speed 400 Hz and a resolution of 1024 x 1024 pixels or 2048 x 2048 pixels. Image analysis was performed using Volocity 6 or 6.3 image processing software (Perkin Elmer).

### Clonal lineage tracing

After immunostaining wholemounts, clones were imaged by confocal microscopy and the number of basal and suprabasal cells in each clone counted in live acquisition mode.

### Primary keratinocyte 3D culture

Mouse oesophagus with the muscle removed were cut into small pieces (2 mm^2^) and placed on a transparent ThinCert^TM^ insert (Grenier Bio-One) with the epitheliem facing upward and the submucosa stretched over the membrane. Inserts in the culture plates were then placed at 37°C for at least 30 min to ensure the attachment of oesophagus explants to the membrane followed by culture in complete FAD medium (50:50) 4.5g/L D-Glucose, Pyruvate, L-Glutamine D MEM (Thermo Fisher Scientific, 11995-065): D-MEM/F12 (Thermo Fisher Scientific, 31330-032), supplemented with 5 µg/ml insulin (Merck, I5500), 1.8×10^-4^ M adenine (Merck, A3159), 0.5 µg/ml hydrocortisone (Merck, 386698), 1×10^-10^ M Cholera toxin (Merck, C8052), 10 ng/ml Epidermal Growth Factor (EGF, PeproTech EC Ltd, 100-15), 5% feral calf serum (PAA Laboratories, A15-041), 5% Penicillin-Streptomycin (Merck, P0781), 1% Amphotericin B (Merck, A2942) and 5 µg/ml Apo-Transferrine (Merck, T2036). Explants were removed after 7 days once half of the membrane had been covered with keratinocytes and the culture was maintained by changing media every three days. When culture established the confluent monolayer, Cholera toxin, EGF and hydrocortisone was removed from the medium to promote differentiation and stratification resulting in multi-layered keratinocyte sheet.

### Virus transduction

To induce cre-recombination in vitro, cultured keratinocytes were infected with Adenovirus encoding Cre recombinase (Ad-CMV-iCre, Vectorbiolabs, #1045 UK). Briefly confluent keratinocyte culture was dissociated by trypsinization, resuspended in adenovirus-containing medium supplemented with polybrene and seeded in the insert. Following 24 hours incubation at 37°C, cells were washed and then cultured in complete FAD medium until confluent. For the experiments, cells were maintained in FAD media without cholera toxin, EGF and hydrocortisone. For non-recombined control, similarly cells were infected with empty Adenovirus.

### *In vitro* cell competition assay

Esophageal keratinocytes were isolated from *Trp53^flR245W-GFP/wt^* and *Trp53^flR245W-GFP/-^* mice and cultured in 3D and *p53\** mutation was induced fully by adenovirus infection as described above. As *p53^wt/wt^* and *p53^wt/-^*, same set of cells infected with empty adenovirus were used. Established cultures were trypsinized, mixed 1:3 (*p53^*/wt^* or *p53^*/-^* : *p53^wt/wt^* or *p53^wt/-^*) and seeded in complete FAD medium. When cultures are fully confluent, cholera toxin, EGF and hydrocortisone was removed from the medium (day 0) and maintained for 28 days. Proportion of GFP+ *p53^*/wt^* or *p53^*/-^* cells in culture was determined by flow cytometry at day 0 and day 28, and data were presented as fold change. Duplicate cultures were immunostained with anti-GFP as described above and imaged by confocal microscopy.

### Flow cytometry

Flow cytometry was carried out using CytoFLEX (Beckman Coulter Life Sciences) flow cytometer. GFP fluorescence was collected using the 488 nm laser and a 525/40 bandpass filter. Data was analysed using FlowJo v10.6.1.

### RNA-Seq and expression analysis

Total RNA was extracted from 3D cultures of mouse primary keratinocytes using RNeasy Micro Kit (QIAGEN, UK), following the manufacturer’s instructions, including DNase digestion. For RNAseq, libraries were prepared in an automated fashion using an Agilent Bravo robot with a KAPA Standard mRNA-Seq Kit (KAPA BIOSYSTEMS). In house adaptors were ligated to 100-300 bp fragments of dsDNA. All the samples were then subject to 10 PCR cycles using the sanger_168 tag set of primers and paired-end sequencing was performed on Illumina’s HiSeq 2500 with 75 bp read length.

Reads were aligned using STAR (v2.5.3a) to the GRCm38 mouse genome Alignment files were sorted and duplicate-marked using Biobambam2 (v2.0.54). Read counting was performed using the script htseq-count (v0.6.1p1) ^36, 37^ and differential gene expression was analyzed using DESeq2 (v1.2.0)^38^. Downstream analysis and visualization was performed using R (v3.5.1, http://www.R-project.org/) with the packages pheatmap (v1.0.10, https://CRAN.R-project.org/package=pheatmap), clusterProfiler (v3.8.1, http://www.bioconductor.org/packages/devel/bioc/html/clusterProfiler.html)^39^ with mouse annotation provided by org.Mm.eg.db (v3.6.0, http://bioconductor.org/packages/org.Mm.eg.db/). Heatmaps were generated from the Transcript Per Million (TPM) values using Morpheus (https://software.broadinstitute.org/morpheus/).

### DNA sequencing

#### 2 mm^2^ gridded samples

Esophageal epithelium prepared as described above was cut into a contiguous array of 2mm^2^ biopsies. DNA was extracted from samples using QIAMP DNA micro kit (Qiagen, catalog no. 56304) following manufacturer’s instructions. DNA from the ears of the same mice was also extracted in the same way and used as germline controls.

#### Low input sequencing samples

Esophageal epithelium was stained for GFP to identify *p53* mutant areas and 0.25 mm diameter samples were taken from GFP positive, GFP negative and microscopic lesions using a brain punch biopsy (Stoelting Europe) under fluorescent dissection microscope. Frozen sections of macroscopic tumors were stained with H&E and area of interest was scraped off from the slide using scalpel. Samples were extracted using the Arcturus Picopure Kit (Applied Biosystems) following the manufacturer’s instructions.

### Sequencing and alignment

Targeted-enriched samples were multiplexed and sequenced on the Illumina HiSeq 2000 v4 platform using paired-end 75-base pair reads. WGS was also performed on HiSeq 2000 v4 platform using paired-end 125-base pair reads. Paired-end reads were aligned to the GRCm38 reference with BWA-MEM (v.0.7.16a & v0.7.17, https://github.com/lh3/bwa). Samples were collected and sequenced over 2 years during which BWA-MEM was updated from v0.7.16a to v0.7.17. As noted in the release notes for v0.7.17 this release adds features but did not alter the alignment output if parameters were not changed, hence remapping is not required for the re-run v0.7.16a data sets. Optical and PCR duplicates were marked using Biobambam2 (v.2.0.86, https://gitlab.com/german.tischler/biobambam2). Samples were mapped to the GRCm38 reference. Mean depth of coverage for each set of sequencing can be found in Supplemental tables 12, 20, 22 and 24.

### Custom baitset design

A custom bait capture to target the exonic sequences of 142 genes was designed using Agilent SureDesign. The list of genes selected for ultra-deep targeted sequencing is shown below:

*abcb11, abcc2, adam10, aff3, ajuba, akt1, apob, arid1a, arid2, arid5b, asxl1, atm, atp2a2, atrx, b2m, bcl11b, braf, brca2, cacna1d, card11, casp8, ccnd1, cdh1, cdkn2a, cobll1, col12a1, crebbp, csmd2, ctcf, ctnnb1, ctnnd1, cul3, cyld, dclk1, dclre1a, ddr2, dicer1, dll1, dll3, dnm2, dnmt3a, dst, dtx1, dtx3, egfr, eif2d, ep300, epha2, erbb2, erbb3, erbb4, ezh2, fat1, fat2, fat3, fat4, fbxo21, fbxw7, fgfr1, fgfr2, fgfr3, flt3, grin2a, hmcn1, hras, huwe1, hydin, itch, jag1, jag2, kdm6a, kdr, keap1, kit, klrc3, kmt2c, kmt2d, kras, krtap4-9, lfng, lrp1b, lrp2, maml1, maml2, maml3, met, mfng, mib1, mtor, nf1, nf2, nfe2l2, notch1, notch2, notch3, notch4, nras, nsd1, numb, numbl, pik3ca, prex2, psen1, psen2, psme4, ptch1, pten, rac1, rasa1, rb1, rbpj, rfng, rhbdf2, rita1, robo1, robo2, ros1, rpgrip1, rpl10, ryr2, scn10a, setd2, slit2, smad4, smarca4, smo, sox2, spen, stat5b, sufu, tert, tet2, tgfbr1, tgfbr2, trp53, trp63, tsc1, vhl, vmp1, zan, zfhx3, zfp750.*

### Variant Calling

#### ShearwaterML

Somatic mutations are generally identified by detecting mismatches between samples (typically comparing tumour with normal samples sequenced at relatively low coverage of 20-40x). Here, in order to identify somatic mutations present in a small fraction of cells within the samples we used the ShearwaterML algorithm from the deepSNV package (v1.21.3, https://github.com/gerstung-lab/deepSNV) to call for mutation events on ultra-deep targeted data ^24, 27, 40^. Instead of using a single-matched normal sample, the ShearwaterML algorithm uses a collection of deeply-sequenced normal samples as a reference for variant calling that enables the identification of mutations at very low allele frequencies. For the normal panel, we used 28 germline samples from ears of the mice analysed in this study and additional mice. Summed mean depth of coverage was 6848x and 4394x for 2mm^2^ gridded samples and 0.25 mm diameter punch samples, respectively.

#### CaVEMan and Pindel

For sequencing data of 0.25mm diameter samples, variants were called using the CaVEMan (v1.13.14 & v1.14.0) and Pindel (v3.3.0) algorithms^41, 42^. For SNVs, CaVEMan was run with the major copy number set to 10 and the minor copy number set to 2. Only SNVs that passed all CaVEMan filters were retained. CaVEMan v1.14.0 was released mid-project and was utilised as it offered additional error checking and speed improvements without altering the results from v1.13.14. Additional filtering to remove mapping artefacts associated with BWA-MEM were: the median alignment score of reads supporting a variant had to be at least 140 and the number of clipped reads equal to zero. In addition, the proportion of mutant reads present in the matched sample also had to be zero. Variants with at least one mutant read present in the matched sample were also removed. Two SNVs called at adjacent positions within the same sample were merged to form a doublet-base substitution if at least 90% of the mapped DNA reads containing at least one of the SNV pair contained both SNVs. Small (<200 bp) insertions and deletions were called using Pindel. Only indels that passed all Pindel filters were kept.

Variants were annotated using VAGrENT (v3.7.0) ^43^. Full lists of called and pass-filter variants are in Supplementary Tables 12, 20, 22 and 24.

#### Merging of mutant clones from contiguous samples

In order to avoid counting the same mutation multiple times and to obtain a more accurate estimate of clone sizes, clonal mutations that spanned between two or more adjacent biopsies were merged and considered as singular events. To do this we calculated the mean number of shared mutations between biopsies at increasing distances, since the immediately adjacent samples are predicted to have more shared mutations than distant samples. We decided to merge mutations common between samples closer than 3mm, as the number of shared mutations plateau at distances greater than this.

#### Mutation burden

The average number of mutations per cell in a given sample was estimated from the VAFs and the number of bases within the bait set with synonymous mutations (dubbed as synonymous footprint), as described previously^24^. When using targeted sequenced data, the mutation burden can be inflated by the alterations in strongly selected genes. Therefore, only the synonymous mutations were used for this calculation.

#### Clone sizes and coverage

The size of mutant clones within each sample can be calculated from the area of the biopsy (2mm^2^) and the fraction of cells carrying a mutation within the sample, as described previously^24, 27^. The lower (=VAF) and upper (=2xVAF) bound estimates of the percentage of epithelium covered by clones carrying non-synonymous mutations in a given gene was calculated for each biopsy, this range allows for uncertainty in copy number. The fraction of epithelium covered by the mutant genes was then calculated from the mean of summed VAF (capped at 1.0) of all the biopsies in the same tissue.

### Gene selection (dN/dS)

We used the maximum-likelihood implementation of the dNdScv algorithm (v0.0.1.0, https://github.com/im3sanger/dndscv) to identify genes under positive selection^24^. dNdScv estimates the ratio of non-synonymous to synonymous mutations across genes, controlling for the sequence composition of the gene and the mutational signatures, using trinucleotide context-dependent substitution matrices to avoid common mutation biases affecting dN/dS. Values of dN/dS significantly higher than 1 indicate an excess of nonsynonymous mutations in that particular gene and therefore imply positive selection, whereas dN/dS values significantly lower than 1 suggest negative selection. In our experimental set up, only mutations that reach a minimum size are detected by deepSNV. Therefore, a significant value of dN/dS>1 indicates that a clone that acquires a non-synonymous mutation in that particular gene will have a higher probability to reach a detectable clone size as compared to a synonymous mutation in the same gene. Hence, genes with dN/dS>1 are considered drivers of clonal expansion.

### Mutational spectra and signature

Mutational spectra for single base substitutions were plotted and compared to 65 known mutational signatures using linear decomposition with the deconstructSigs R package (v1.9.0, https://github.com/raerose01/deconstructSigs)^44^. The mutational spectra of DEN-treated samples were highly consistent, precluding deconvolution into separate signatures (either known or de novo). Transcriptional strand bias was analysed using MutationalPatterns (v3.4.0, https://bioconductor.org/packages/release/bioc/html/MutationalPatterns.html)^45^.

### Copy number analysis

The copy number analysis workflow was conceptually very similar to that used in ^24^. That paper described analysis of whole genome amplified exome sequenced triples. Here we utilise whole genome and targeted sequencing, which required adaptation of the workflow. For clarity and completeness, the workflow is described in full here.

#### Whole genome sequencing

A modified version of QDNAseq (https://github.com/ccagc/QDNAseq/) was used to call changes in total copy number from the low coverage (mean coverage 1.8x) whole genome sequencing data^46^. QDNAseq was modified to include the correction of the coverage profile of the sample of interest by that of a matched control. The procedure to call gains and losses is as follows: First sequencing reads were counted per 100kb bins for both the sample of interest and the matched control. The bin-counts were then combined into coverage log ratio values to obtain what is commonly referred to as “logR”. The calculation of logr is implemented similarly to how the Battenberg copy number caller calculates these values^47^: first the bin-counts from the sample of interest were divided by the control bin-counts to obtain the coverage ratio; the coverage ratio was then divided by the mean coverage ratio and finally the log2 was taken to obtain logr. The regular QDNAseq pipeline is then applied: First a correction of the logr for GC content correlated wave artifacts, then segmentation and finally calling of gains and losses.

#### Targeted sequencing

The workflow to analyse targeted sequencing data differs slightly from the above whole genome workflow. When performing targeted sequencing, only a subset of the genome is actively targeted and yields high coverage. The remainder of the genome is typically covered by much shallower, off-target coverage^48^. Removal of the targeted regions (typically a small proportion of the genome) effectively yields a shallow whole genome sequencing sample.

The read counting step was therefore adapted to only count off-target reads. We first counted all reads per 1Kb bins along the genome, then removed any bin that overlaps with the start/end coordinates of a region contained within the kit and mapped the remaining onto bins of 1Mb. This procedure was applied to both the sample of interest and the matched control, after which the construction of logr normalises out the fact that some bins span slightly less than 1Mb due to removal of the targeted regions. The above-described pipeline is subsequently run to obtain copy number calls.

A comparison of calls between samples from whole genome and targeted sequencing from the same mice shows good correspondence of the overall picture gained from copy number analysis (**Fig. 4h and Extended Data Fig. 5f**).

#### Post-hoc filtering of calls

A post-hoc filtering step was subsequently applied to obtain robust copy number calls. The calls from both whole genome and targeted sequencing were required to constitute an alteration in at least 10% of sequenced cells and be at least 20Mb in size.

### Code availability

The pipeline code and modified QDNAseq package are available at https://github.com/sdentro/qdnaseq_pipeline and https://github.com/sdentro/QDNAseq/tree/dev respectively.

### Data availability

The sequencing data sets in this study are available at the European Nucleotide archive (ENA) Accession numbers for RNAseq data on http://www.ebi.ac.uk/ena are as follows: *p53^wt/wt^*, ERS1432686, ERS1432689, ERS1432688, ERS1432703, ERS1432704, ERS1432705; *p53^*/wt^* , ERS1432636, ERS1432637, ERS1432634, ERS1432638, ERS1432635, ERS1432639; *p53^wt/-^*, ERS1432650, ERS1432651, ERS1432656, ERS1432654, ERS1432657, ERS1432658; *p53^*/-^*, ERS1432672, ERS1432670, ERS1432671, ERS1432673, ERS1432674, ERS1432676.

Accession numbers for targeted DNA sequencing samples and Whole genome sequencing are ERP129331 and ERP129332 respectively.

### TCGA analysis

The TCGA ESCA project dataset contains the largest publicly available complete dataset with SNV, CNV and LOH calls from whole genome sequencing of ESCC samples ^49, 50^. Sample metadata was obtained from the NIH GDC data portal (https://portal.gdc.cancer.gov/). Within the ESCA project filters were applied to select only squamous cell carcinoma samples with a matched blood normal sample. Application of filters resulted in 88 samples remaining for analysis. Somatic SNV and CNV calls were also obtained from the GDC portal. LOH calls for TCGA were published previously and are also available from the GDC pancan portal (https://gdc.cancer.gov/about-data/publications/pancan-aneuploidy)^49^. Median age across the ESCC sample cohort was 57, atypical of the incidence in Europe which peaks in the age group 85 to 89 for females and above 90 for males (http://www.cancerresearchuk.org/health-professional/cancer-statistics/statistics-by-cancer-type/oesophageal-cancer/incidence.)

Variants overlapping *TP53* were extracted for each of the samples and were summarised in **Supplementary table1**. For each sample the most severe SNV consequence in *TP53* was identified in the order: Dominant negative, nonsense, missense, other & none. Identified dominant negative mutations defined as the following previously identified in literature: p.R175H, p.Y220C, p.M237L, p.R248Q, p.R248W, p.R273H, p.R282W & p.R249S ^32, 51^.

### Statistical analysis

Data are expressed as mean values ± s.e.m. Statistical analysis was performed using the Graphpad Prism software and can be found in Supplementary Tables. No statistical method was used to predetermine sample size. The experiments were not randomized. The investigators were not blinded to allocation during experiments or outcome assessment.

### Spatial simulations of *p53*^*/wt^ mutant clone growth in mouse oesophagus

*p53*^*/wt^ mutant cell behavior was modelled as a non-neutral single progenitor (SP) model with a fate imbalance (*δ*) in the mutant population^15^. This imbalance leads to an increased probability of symmetric divisions producing two mutant progenitors, effectively providing a fitness advantage over wild-type cells.

Here we adapted the spatial single progenitor model described in Kostiou et al implemented in NetLogo^52, 53^. Briefly, we simulate the imbalanced single progenitor as a two-dimensional stochastic cellular automaton (CA) model in a hexagonal lattice (**Extended Data Fig. 2a**). Basal cells were simulated on a lattice of 10,000 cells (1% of the area of adult mouse oesophagus), with periodic boundary conditions. Each simulation was repeated 100 times. To take account of local crowding events lattice site may be occupied by either by no cell, single cells, or pairs of cells. Doubly occupied sites transition to single occupancy once immediately adjacent empty sites become available or through an instant “extrusion” event, if one cell is committed to the differentiated fate. Based on previous observations of *p53*^*/wt^ mutant cell growth in mouse skin, mutant cells were set to respond to adjacent double-occupancy sites by reducing their propensity to symmetric division giving dividing cells (**Fig. 1b,2f**)^53^.

#### Model parameter estimation

To estimate the key parameter representing fate imbalance, following experimental observations that *p*53**^*/wt^** cells divide and stratify at the same rates as their wild type counterparts, we made the simplifying assumption that other parameters remain unchanged in the mutant population. This can be estimated from a linear regression applied to the *ln(average clone size)* ^54^, where the slope of the line is a function of the known parameters and the imbalance.

We used parameter values describing the homeostatic cell population dynamics in mouse oesophageal epithelium provided by Doupe et al ^15^. To test the robustness of these parameters, two more parameter combinations are were taken from different experiments the mouse oesophagus^16^. All available parameter sets proposed for this tissue were simulated (**Extended Data Fig. 2**). The parameter combinations and resultant imbalance (δ) were the following:

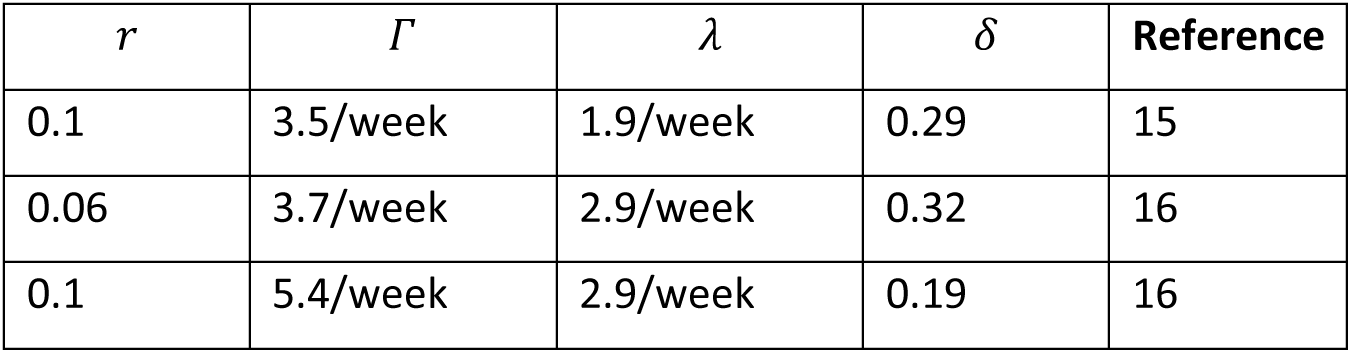

| $r$ | $\Gamma$ | $\lambda$ | $\delta$ | Reference |
| --- | --- | --- | --- | --- |
| 0.1 | 3.5/week | 1.9/week | 0.29 | 15 |
| 0.06 | 3.7/week | 2.9/week | 0.32 | 16 |
| 0.1 | 5.4/week | 2.9/week | 0.19 | 16 |

We further tested the robustness of the parameter estimates by perturbing estimates of *λ* and *r*, find that there was a broad agreement between simulations and observations (**Extended Data Fig. 2b,c**).

Analysis of the simulated p53 mutant cell populations in the basal layer of mouse oesophagus demonstrates that the model is in quantitative agreement with the experimental values and is broadly robust to the different parameters proposed for this tissue (**Extended data Figure 2**). This confirms that a spatial non-neutral single progenitor model with cell density dependent feedbacks is able to recapitulate the observed p53 mutant behaviour in the mouse oesophageal epithelium.

---

**Extended Data Fig. 1:**
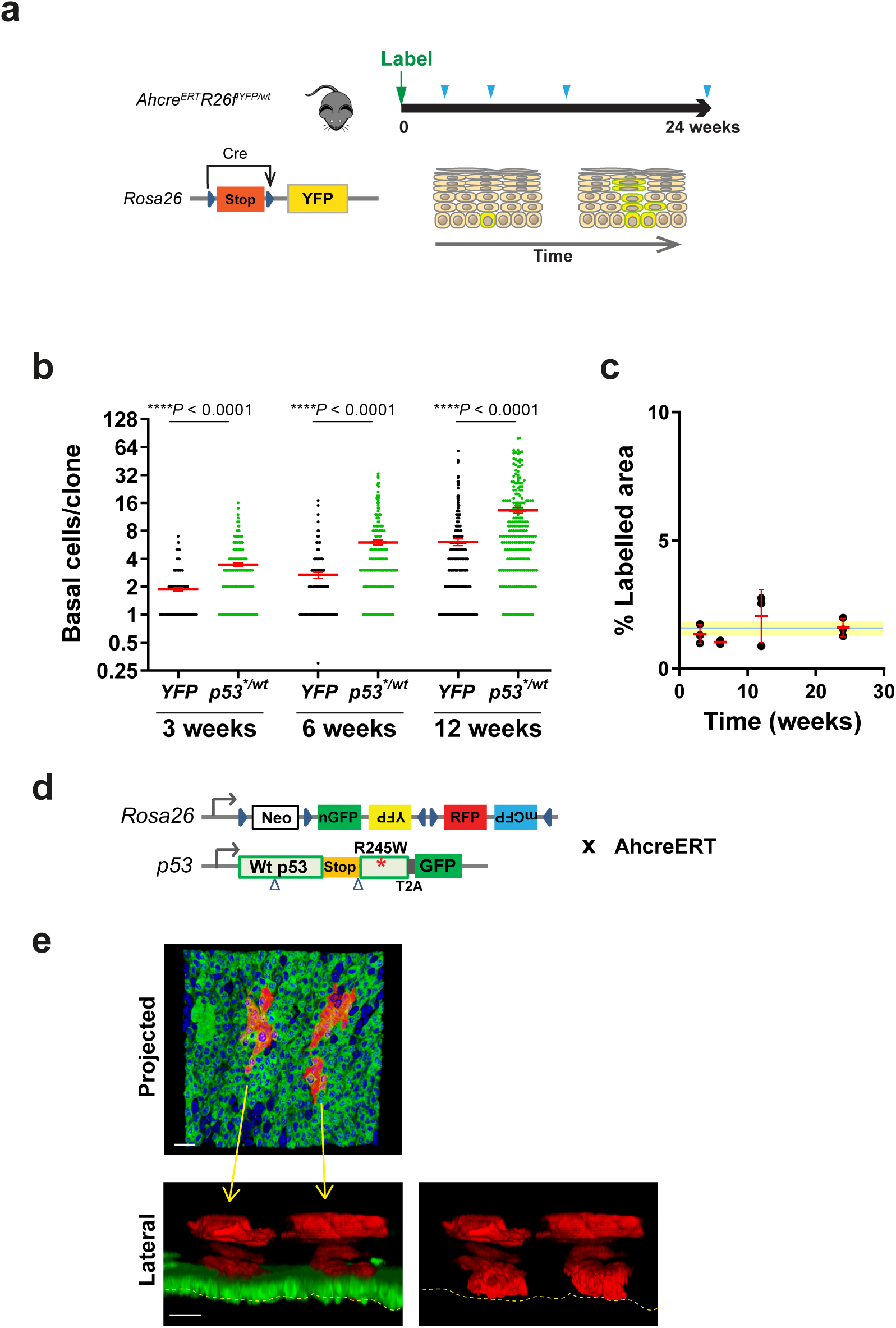
Behaviour of wild type and p53 mutant clones in homeostatic mouse esophagus. **a,** Lineage tracing protocol: Clonal frequency labelling was induced in the esophageal epithelium of adult *Ahcre^ERT^ R26^flYFP/wt^* (*p53^*/^*^wt^) mice and samples were collected at indicated time points (blue arrowhead). **b,** Comparison of basal clone size. Clone size distribution of *p53^wt/wt^* (YFP) and *p53^*/^*wt (GFP) (for ≥2 cell clones). Error bars indicate mean ± s.e.m; p values by Mann Whitney test. 2-4 mice per time point. n=189 YFP and 247 GFP clones at 3 weeks, 140 YFP and 230 GFP clones at 6 weeks, 224 YFP and 223 GFP clones at 12 weeks. **c,** Projected labelled area of YFP clones. Error bars indicate mean ± s.e.m. Blue line and shading show average and s.e.m. across all time points. n=11 mice. See **Supplementary Tables 2 and 3**. **d,** Multicolor lineage tracing in *Ahcre^ERT^ Rosa26^flConfetti/wt^ p53^*/wt^* animals. The fate of *p53* mutant progenitors can be tracked in double labelled clones even if the p53 locus becomes inactive in the differentiated progeny. **e,** Rendered z stacks of typical *Confetti (RFP)-p53^*/wt^* double labelled clone12 weeks post induction of *Ahcre^ERT^ Rosa26^flConfetti/wt^ p53^*/wt^* animals. Due to the high level of induction, clones were fused by 12 week time point. However double labelled clones could be identified as shown in this image. Red, RFP, Green, GFP, Blue, DAPI. Scale bar, 20 µm.

**Extended Data Fig. 2:**
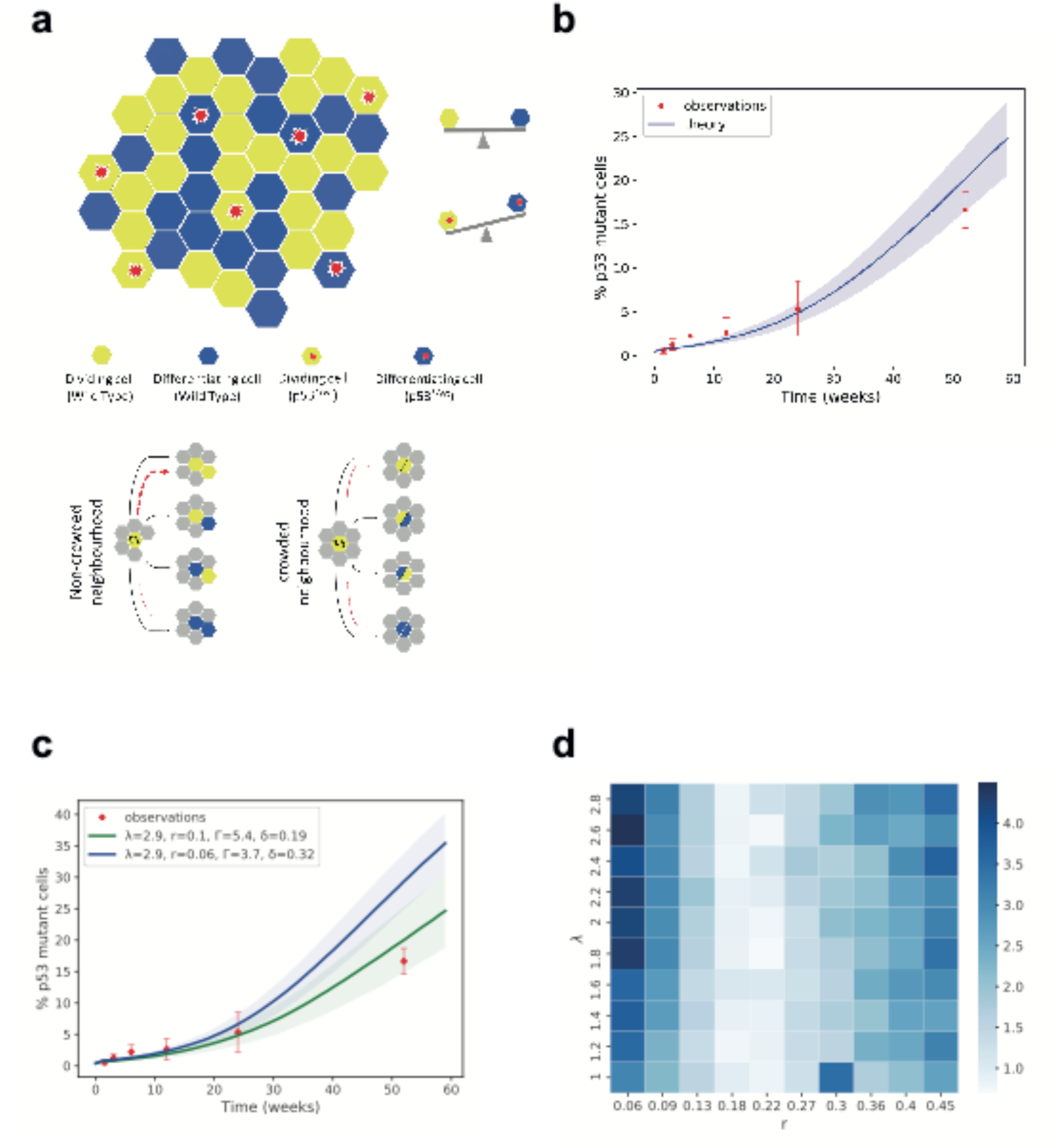
Model of p53^*/wt^ mutant clone dynamics. The growth of *p53* mutations in the basal layer of the mouse oesophageal epithelium is successfully reproduced by a spatial non-neutral SP model with a cell density dependent feedback that mutant cells respond to **a**: Upper panel: Illustration of the two-dimensional hexagonal lattice representing the basal layer. Proliferating cells are shown in yellow and differentiating cells in blue. Proliferating cells divide, producing two daughter cells, whilst differentiating cells exit the basal layer and are removed from the simulation. Mutant cells, marked with a red asterisk, have a bias towards producing proliferating daughters. Lower panel: Schematic representation of the rules of the spatial model. Proliferating cells undergo a division type with balanced division probabilities. In case of mutant cells, division probabilities are biased, favouring symmetric division, indicated by dashed red lines. Mutants lose this advantage in response to crowding in their local neighbourhood and switch their behaviour to balanced dynamics. **b**: Tissue colonisation plot showing the simulated (solid lines) and observed (red points) percentage of mutant cells over time. Simulated data correspond to mean values across 100 simulations. Shaded area and error bars correspond to SD. Parameters used: *r* = 0.1, *Γ* = 3.5/*week*, *λ* = 1.9/*week*, from^15^. Fate bias was calculated as *δ* = 0.29, from a linear regression applied to the *In(average clone size)*, where 2*δrλ* = *slope*. Crowding threshold was set to 7 cells, see methods for details. **c**: Analysis of the spatial non-neutral SP model with cell density feedback, simulating p53 mutant growth in the basal layer of mouse oesophageal epithelium under different inferred parameter sets (see methods). Plot showing the simulated (solid lines) and observed (red points) percentage of mutant cells over time. Simulated data correspond to mean values across 100 simulations. Shaded areas and error bars correspond to SD. The values of *r*, *Γ*, *λ* were taken from^16^. The value of fate bias (*δ*) was calculated from a linear regression applied to the *In(average clone size)*, where 2*δrλ* = *slope*. **d**: Exploration of spatial model’s fit sensitivity to different parameter combination regimes (see methods). Heatmap shows the root mean squared error (*RMSE*) as measured from the distances between the simulated and observed *p53* mutant cell percentages. Simulated mutant cell percentages correspond to mean values across 100 simulations. Spatial simulations were performed for 10×10 *r*, *λ* parameter combinations, with 100 repetitions for each combination. Fate bias (*δ*) values for each combination were calculated from a linear regression applied to the *In(average clone size)*, where 2*δrλ* = *slope* and assuming that the *In(average clone size)* of the simulated data would have the same slope as the observed data.

**Extended Data Fig. 3:**
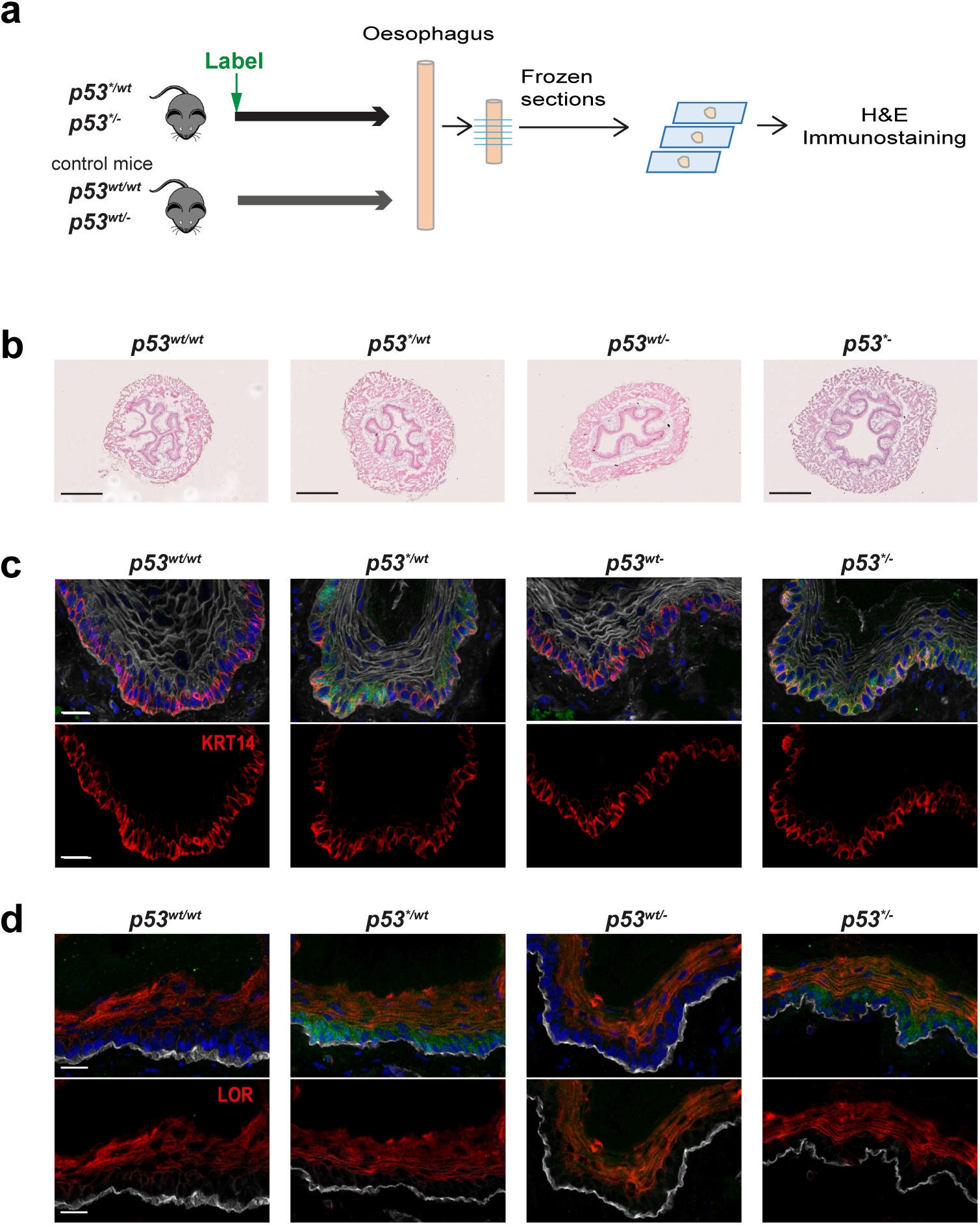
Effect of *p53\** mutation on esophageal epithelium. **a,** Protocol: Cryosections of mouse esophageal epithelium from mice with indicated genotypes were generated as shown, *p53^*/wt^* and *p53^*/-^* esophagus was harvested at 8 months post high level induction. Representative images from 3 mice per group. **b,** H and E. Scale bars, 500µm. **c,** Basal layer, KRT14 (red). Green, GFP; grey, WGA. Scale bar, 20µm. **d,** Differentiated cells in suprabasal layer, LOR (red). Green, GFP; grey, ITGA6. Scale bar, 20µm.

**Extended Data Fig. 4:**
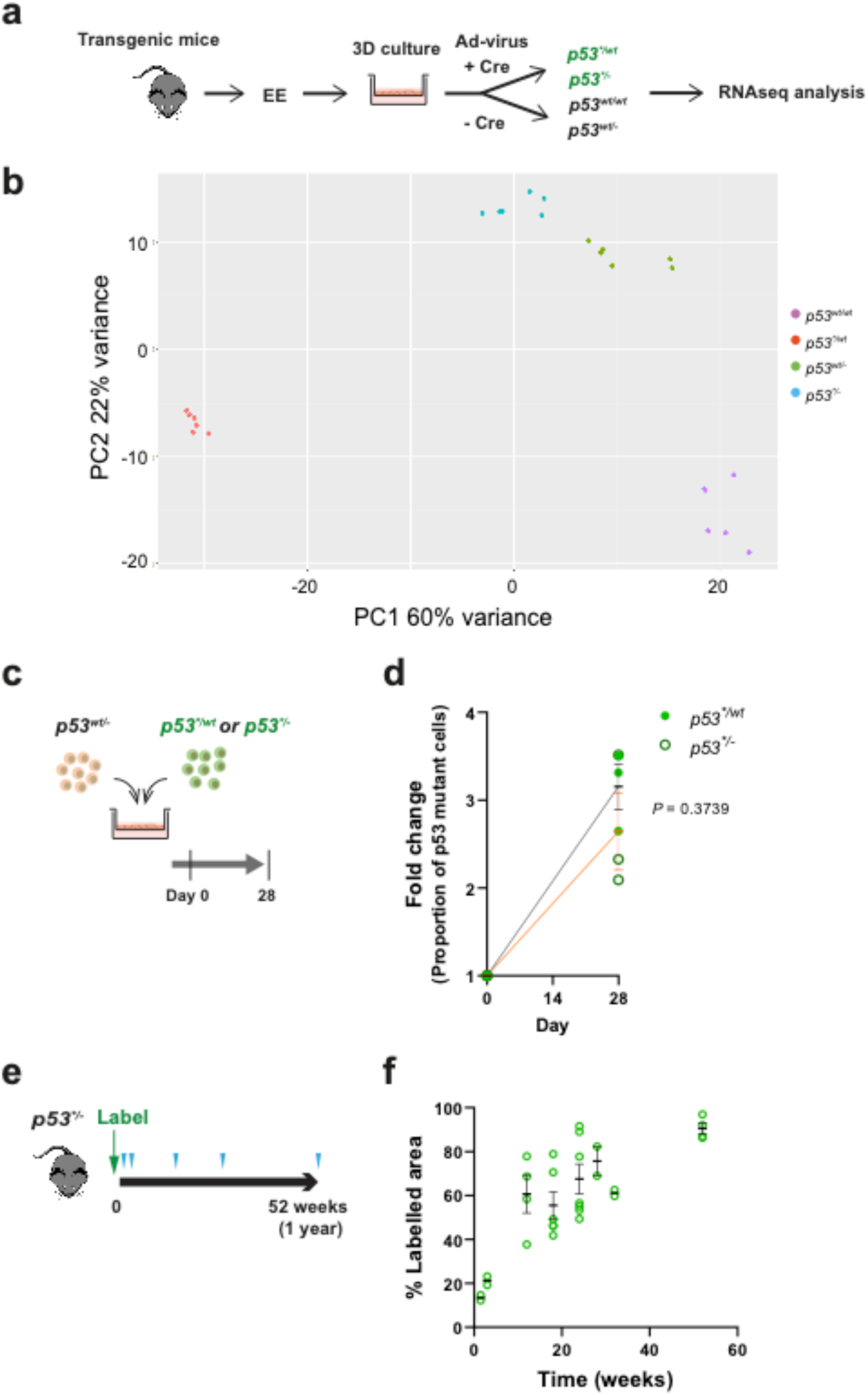
Characterization of *p53* mutant cells. **a,** Primary esophageal keratinocytes from transgenic mice were culture in 3D and *p53\** mutation was induced using adenovirus encoding Cre recombinase. **b,** Principal component analysis of RNAseq data showing the variation of gene expression profile between samples. Total RNA was collected from the primary cultured esophageal keratinocytes of indicated genotype and RNAseq analysis was carried out. **c,** Protocol: *p53^*/wt^* and *p53^*/-^* cells were cocultured with *p53^wt/-^* cells respectively and relative fitness was examined. **d,** Quantitation of cell competition assay by flow cytometry. Graph shows the fold change of proportion of GFP+ *p53^*/wt^* or *p53^*/-^* cell in the culture. Black (*p53^*/wt^*) and red (*p53^*/-^*) lines indicate mean and s.e.m. *P* value was determined by two-tailed Student’s t-test. **e,** *in vivo* analysis of *p53*/-* clone behaviour. Expression of *p53* mutant and GFP reporter was induced clonally (Labelling) and esophagus samples were taken at indicated time points (triangles). The fate of mutant clones was examined by tracking the expression of GFP. **f,** Proportion of labelled projected area at indicated time points. Error bars are mean ± s.e.m. n=2 mice for 1.5, 3, 28 and 32 week time points, n=4 for 12 week, n=6 for 18 week, n=7 for 24 week and n=4 for 52 week time points. See **Supplementary Tables 7-10**.

**Extended Data Fig. 5:**
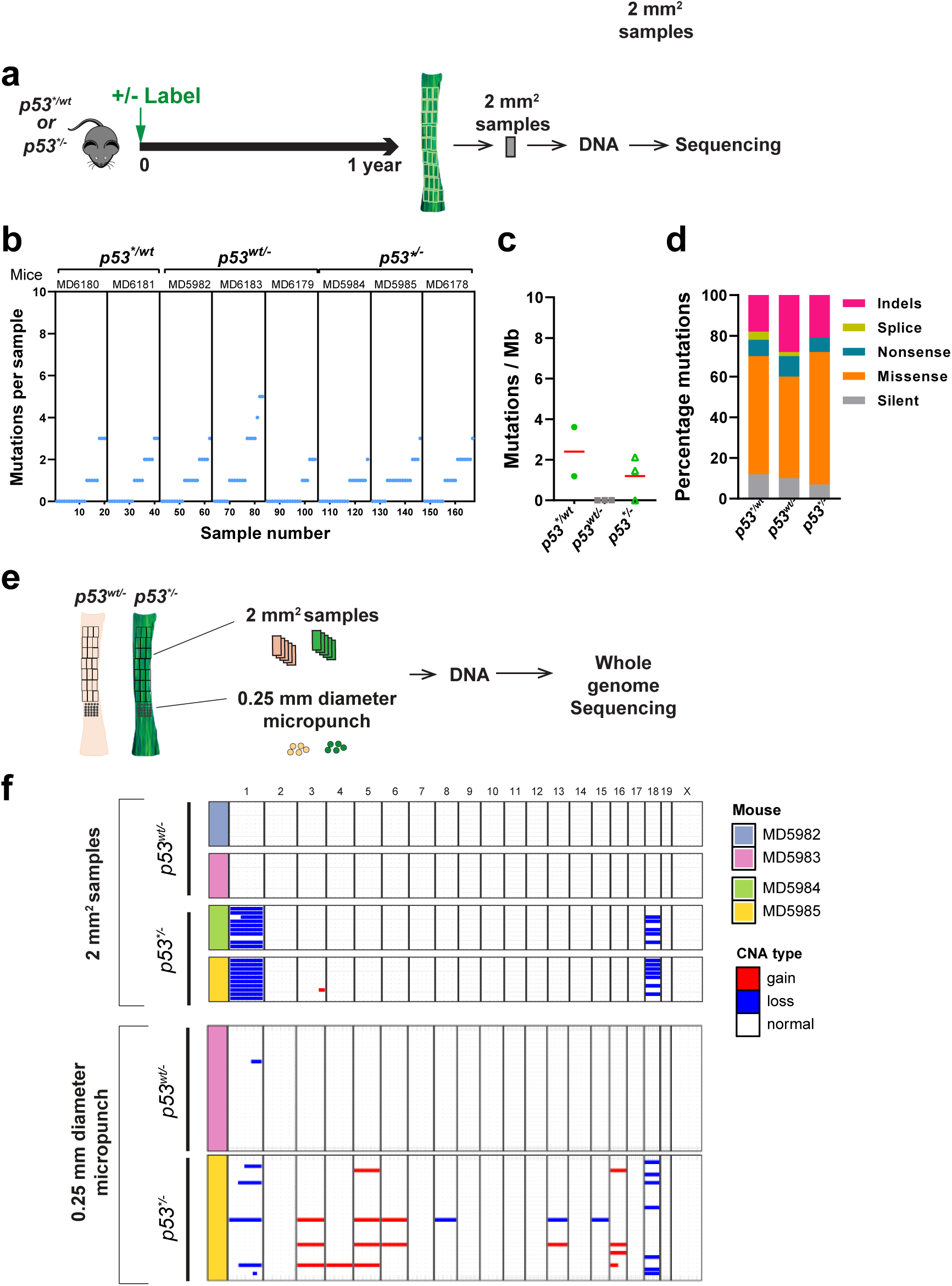
DNA sequencing analysis of *p53* mutant EE. **a,** Protocol: EE from indicated mice were collected at 1 year time point and cut into 2mm^2^ grid pieces. DNA from each sample was analysed by ultradeep targeted sequencing (a-d) or WGS (e). **b,** Number of mutations per sample. Every dot corresponds to a sample. n=2 mice for *p53^*/wt^* and n=3 mice for *p53^wt/-^* and *p53^*/-^***. c,** Estimated mutation burden in indicated genotype. No significant difference between genotypes by unpaired Student’s t-test. Red lines indicate mean value. **d,** Percentage of mutation types identified in each genotype. **e,** Copy number analysis comparing sample size. 2 mm^2^ and micro-punch biopsies were taken from same EE and subjected to whole genome sequencing. **f,** Summary of copy number analysis. The result with 2 mm^2^ samples was comparable to that detected by ultradeep targeted sequencing (Fig. 4h), whereas micro-punch biopsies showed more variety of copy number alterations. See **Supplementary Table 12 and 13**.

**Extended Data Fig. 6:**
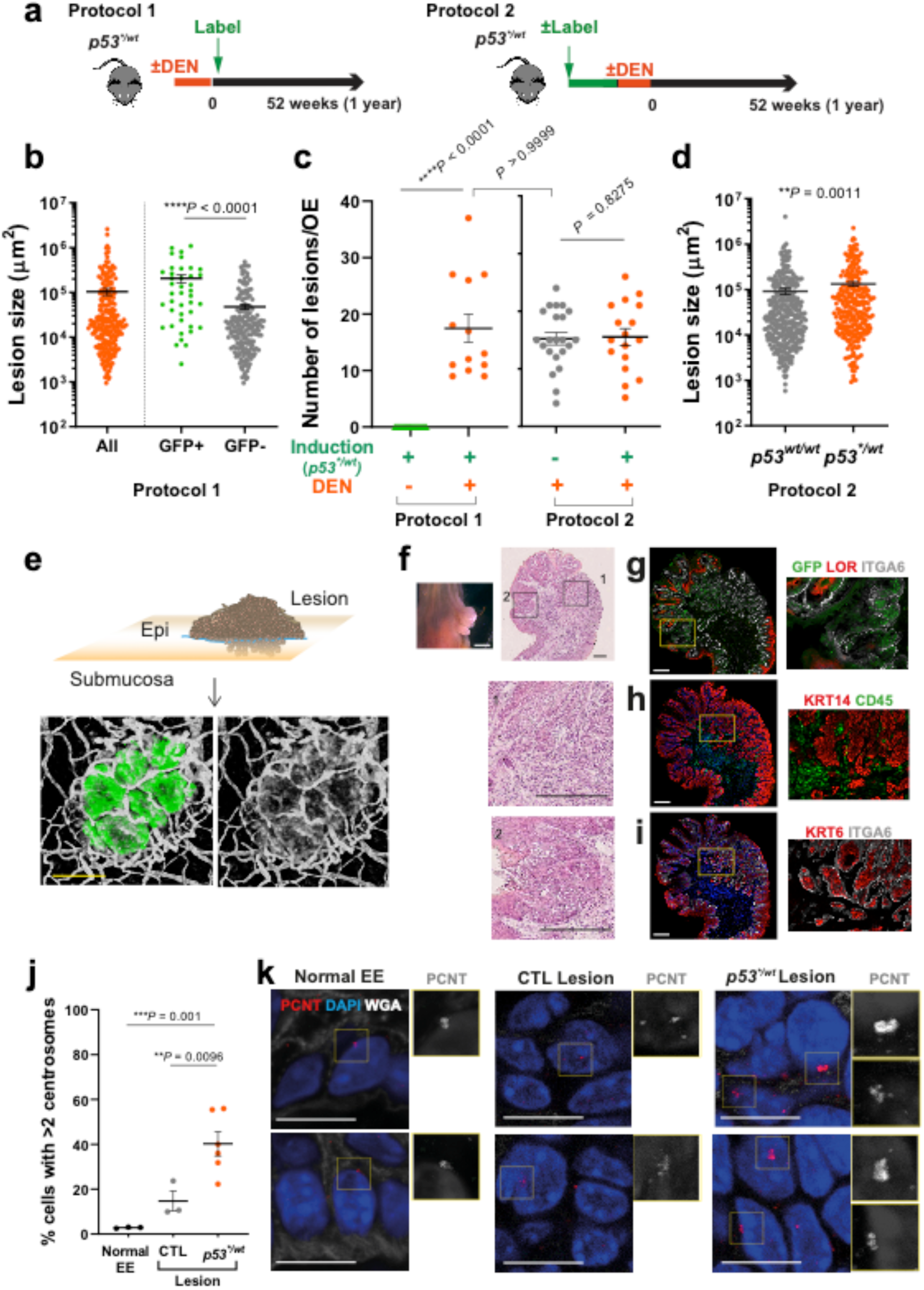
Lesions formed in mutagenized epithelium carrying *p53^*/wt^* clones. **a,** Protcol 1: *p53\** mutation was induced after DEN treatment, *p53^*/wt^* clones competed in mutated EE, less than 10% of them persisted. Protocol 2: Induced *p53^*/wt^* mice were treated with DEN, introducing mutations within *p53^*/wt^* cells which had expanded to occupy >70% of the epithelium. Lesions found in these mice after 6 months were quantified. **b,** Comparison of lesion size in protocol 1. Error bars indicate mean ± s.e.m.; two-tailed Mann Whitney test. n=227 lesions from 13 mice. Contribution of *p53^*/wt^* to the lesions was also assessed by GFP expression (GFP+). **c,** Number of lesions. Error bars indicate mean ± s.e.m. n=22 for induced *p53^*/wt^* control (untreated), n=13 for DEN-treated induced mice, n=22 for uninduced mice (*p53^wt/wt^)* and n=17 for *p53^*/wt^*. *P* values were determined by two-tailed Mann Whitney test. **d,** Comparison of lesion size in protocol 2. The area of lesions were recorded at late time points. Error bars indicate mean ± s.e.m; two-tailed Mann Whitney test. n=335 lesions from 22 mice for *p53^wt/wt^* and n=264 lesions from 17 mice for *p53^*/wt^*. **e,** Schematic of lesion and 3D confocal images of submucosa showing invasion of epithelial cells. Esophageal samples were collected from *p53^*/wt^* induced mice at 6 months post DEN treatment. After the epithelium was peeled off, the submucosa was stained for GFP (*p53^*/wt^*, green), KRT6 (red) and CD31 (grey). The dotted yellow boxes are shown at higher magnification. Scale bars, 200µm. **f,** Image of ESCC and histology. White scale bar 1mm, black scale bars, 200 µm. **g,** GFP for transgenic *p53* mutant expression; LOR, differentiated keratinocyte; ITGA6, basal membrane. **h,** KRT14, progenitors in basal layer; CD45, leukocytes. **i,** KRT6, hyperplastic epithelium marker; ITGA6. Scale bars, 200µm. **j,** Quantitation of centrosome per cell in macroscopic lesions at one year time point. Frozen sections of macroscopic lesions were stained with PCNT (**Extended Data Fig.6k**). Centrosomes were quantified based on number of PCNT foci and size (area). Values are the average number of centrosomes per cell from each normal EE or lesion. Error bars indicate mean ± s.e.m; Welch’s t test. n=299 cells from 3 normal mouse EE, n=332 cells from 3 control (CTL) lesions (n=2mice), n=614 cells from 6 *p53^*/wt^* lesions (n=3 mice). **k,** Representative images of centrosomes in macroscopic lesions. Frozen section of normal EE (untreated), CTL (*p53^wt/wt^*) and *p53^*/wt^* lesions were stained for PCNT. Red, PCNT; glue, DAPI; grey, WGA. Enlarged PCNT foci are shown in greyscale. Scale bars, 10µm. See **Supplementary Table 15-19**.

**Extended Data Fig. 7:**
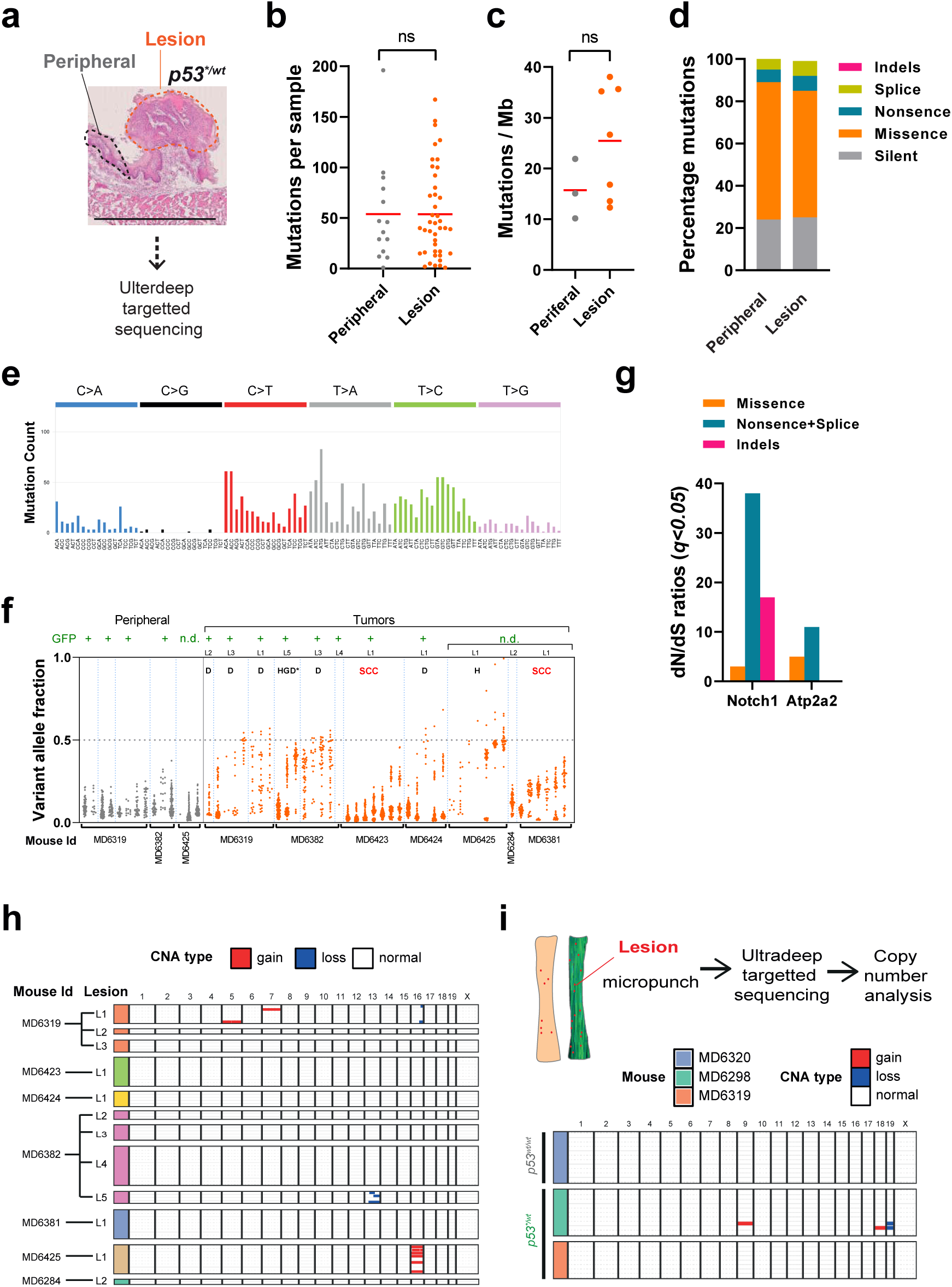
Mutational landscape and CNA in *p53^*/wt^* lesions. **a,** Sequencing samples: macroscopic esophageal lesions from DEN treated *p53^*/wt^* mice were sectioned and stained with H&E. Lesions and peripheral areas were scraped off from slides and DNA was extracted for ultradeep targeted sequencing. **b,** Number of mutations per sample. Each dot corresponds to a sample. Red lines indicate mean value. ns, not significant by unpaired Student’s t-test. **c,** Estimated mutation burden. Each dot represents a mouse. Red lines indicate mean value. ns, not significant by unpaired Student’s t-test. **d,** Percentage of mutation types identified in each group. **e,** Mutational spectrum of lesions. **f,** Variant allele fraction for the mutations detected in samples. Samples from the same lesion were grouped. Pathology result of lesions were indicated in the panel: D, dysplasia; HGD*, high grade dysplasia with potential invasion; H, hyperplasia; SCC, squamous cell carcinoma. GFP expression was determined where possible and indicated at the top of the panel. **g,** Positively selected somatic mutations in tumors. dN/dS ratios for missense, truncating (nonsense + splice) and indels of mutant genes under selection are shown. **h,** CNA in macroscopic tumors. n = 12 tumors from 7 mice. **i,** CNA in microscopic tumors. 18 tumors from 2 induced mice and 11 tumors from uninduced mouse were analysed by off target reads from ultradeep targeted sequencing, see methods. See **Supplementary Tables 20-22**.

